# High immune receptor clonality in melanoma-draining lymph nodes associates with immune dysfunction and poor survival

**DOI:** 10.64898/2026.06.09.729531

**Authors:** Vincent Walter, Johanna Herold, Sophia Aurelie Feuchter, Sophie Thomä, Mara Venohr, Antonio Vogelsberg, Mustafa Kilic, Fiamma Berner, Lena Nanz, Ulrike Leiter-Stöppke, Tobias Sinnberg, Christian M. Schürch, Anja Ulmer, Paul-Christian Bürkner, Lukas Flatz

## Abstract

Tumor-draining lymph nodes (tdLNs) are critical hubs of anti-tumor immunity but are also vulnerable to tumor-mediated immunosuppression. We analyzed T cell and B cell receptor (TCR/BCR) repertoires and transcriptomes from sentinel and non-sentinel lymph nodes of patients with melanoma from a historical pre–immune checkpoint inhibitor cohort (1994-2002) and an independent contemporary validation cohort (2022-2024). Melanoma-positive lymph nodes exhibited increased immune receptor clonality compared with tumor-free nodes. While average clonality showed no consistent association with outcome, the presence of extreme high-clonality outliers in individual lymph nodes was strongly associated with poor melanoma-specific survival. These outliers were characterized by a loss of lymphocyte-related genes and activation markers, an enrichment of melanocytic transcripts, and the suppression of immune signaling pathways, consistent with local immune dysfunction. Increased clonality was confined to lymph nodes and not observed in the peripheral blood. T cell responses to melanocyte differentiation antigens were infrequently shared between lymph nodes and peripheral blood, highlighting immune compartmentalization.

## Introduction

Secondary lymphoid organs provide structured microenvironments in which antigen-presenting cells and adaptive immune cells interact to initiate and shape immune responses. In cancer, tumor-draining lymph nodes (tdLNs) represent central sites of tumor spread and immune regulation, where lymphoid structures can promote either effective immune activation or immunosuppression with major consequences for tumor control (1–5). Indeed, several studies propose that T cells in the tdLNs serve as precursors of tumor-infiltrating T cells (6–8). At the same time lymph nodes (LNs) possess substantial immunologic potential for anti-tumor immunity but are also targets of tumor-mediated immunomodulation, enabling immune evasion (9). This immunomodulation can be actively enforced by node-resident myeloid cells: medullary sinus macrophages that engulf tumor material induce the regulatory cytokine IL-33 and activate regulatory T cells, and IL-33 expression in melanoma sentinel LNs increases with nodal stage and is associated with worse survival (10).

Melanoma is a highly immunogenic tumor characterized by a high mutational burden and robust immune cell infiltration, which has enabled patients to benefit greatly from immune checkpoint inhibitor (ICI) therapies (11). Large-scale surgical removal of LNs, such as complete LN dissection, has not been shown to confer a survival benefit in sentinel LN-positive melanoma (12). Furthermore, in patients with melanoma LN metastases, neoadjuvant and adjuvant ICI therapy result in markedly improved patient outcomes compared with adjuvant treatment alone (13–15), suggesting that the LNs play an important role in tumor control. These findings have prompted an active debate regarding the impact of surgical removal of tdLNs when ICI is available (16).

As ICI therapies harness adaptive tumor-directed immunity, characterization of adaptive immune cell receptors might improve understanding of their effect. During their development, B and T cells undergo receptor recombination, leading to unique complementarity determining region 3 (CDR3) sequences that can be used as molecular barcodes to track their spread across tissues and time. Technical advances have made sequencing of the B cell receptor (BCR) and T cell receptor (TCR) repertoires to track cells or assess the general state of the adaptive immune system feasible (17). Changes in repertoire characteristics, such as immune receptor clonality, which quantifies the expansion of clones within the immune repertoire, can be used to evaluate the overall state and dynamics of the adaptive immune system.

In cancer, most immune repertoire analyses have focused on the effects of ICI therapy on peripheral blood. Peripheral immune receptor clonality has been linked to clinical outcome and toxicity, with high pre-treatment clonality predicting benefit from PD-1 blockade but not CTLA-4 blockade and increased TCR clonality being associated with immune-related adverse events (18–20). However, these observations are largely restricted to peripheral compartments and provide limited insight into immune dynamics within tdLNs.

Less is known about clonal diversity within tdLNs. Reduced TCR β chain (TRB) diversity and clonal expansion in metastatic LNs have been reported in patients with colorectal and breast cancer, with expanded clones often shared between primary tumors and tdLNs (21–23). In melanoma and other solid tumors, shifts toward less specific and less mature BCR repertoires have also been observed in metastatic LNs (24). However, how immune receptor clonality in tdLNs relates to immune competence and patient outcome remains poorly defined.

Here, we leverage a unique collection of sentinel and non-sentinel LNs collected from melanoma patients treated in the pre-ICI era, together with an independent contemporary validation cohort, to define how adaptive immune receptor repertoires in tdLNs relate to melanoma progression and patient survival. By integrating T and B cell receptor sequencing with transcriptomic profiling, and by mapping T cell responses against melanocyte differentiation-antigens (MDAs) across LNs and peripheral blood, we uncover marked immune compartmentalization and identify LN–restricted immune receptor clonality as a feature associated with immune dysfunction and adverse outcome.

## Results

### Effect of melanoma metastases on LN T and B cell receptor clonality

For the first cohort, we recruited 40 melanoma patients who underwent sentinel LN biopsy and elective lymphadenectomy between 1994 and 2002 from our biobank at the University Hospital Tübingen. As patients were treated in the pre-ICI era, they either received no adjuvant treatment or were treated with adjuvant interferon-α. In cases of distant metastases, patients were treated with chemotherapy. Patients were followed up in the context of our melanoma registry for a median of 10.9 years (characteristics in **Supplementary Table 1**). 118 LNs were analyzed and the number of LNs per patient ranged between 1 and 15. A total of 118 LNs met quality criteria for TCR and BCR profiling, while 114 LNs were additionally subjected to transcriptomic profiling. To simplify repertoire analysis, immune receptor repertoires were described using Simpson clonality, a metric ranging from 0 (indicating a completely even distribution of clones) to 1 (representing a monoclonal repertoire). We chose to utilize Bayesian modeling for the immune repertoire analyses in this manuscript because it enabled usage of all data in a cohort with varying numbers of lymph nodes. Model details can be found in the supplementary information and coefficients are given alongside 95% credibility intervals (95%-CIs).

As melanomas are known to modulate the immune system (9, 11), we hypothesized that the spread of melanoma may affect immune repertoire clonality. Indeed, all TCR (TCR α chain (TRA) = 0.126, 95%-CI = 0.017 - 0.241; TRB = 0.18, 95%-CI = 0.036 - 0.329) and immunoglobulin (IG) gene loci (IG heavy chain (IGH) = 0.283, 95%-CI = 0.08 - 0.513; IG κ chain (IGK) = 0.244, 95%-CI = 0.058 - 0.466; IG λ chain (IGL) = 0.197, 95%-CI = 0.014 - 0.401) were found to be more clonal in pathologically melanoma-positive versus -negative LNs (**Figure 1A**). Looking at the individual patient level, however, it became apparent that this effect is due to changes observed in a subset of patients (such as hist15, hist3, hist8, hist29 and hist11), but is not universal (**Figure 1A**, **Figure 2D-E**).

**Figure 1:**
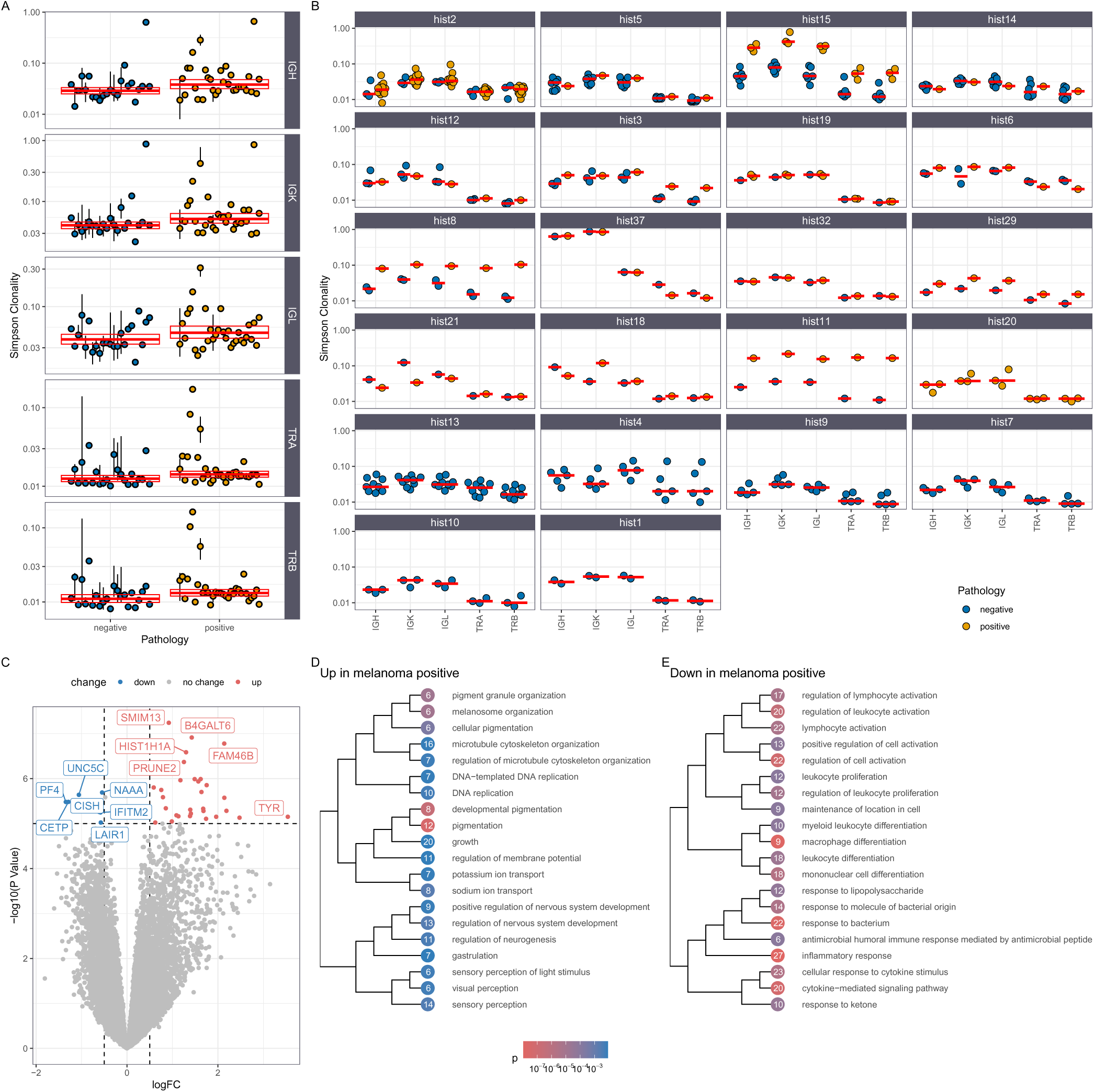
Effect of melanoma spread on lymph node immune repertoire clonality and gene expression. (**A**) Comparison of Simpson clonality in melanoma-positive versus -negative LNs. Circles indicate the patient median and whiskers the intraindividual range. Red lines show the model’s conditional effects with credibility intervals ( IGH = 0.283, 95%-CI = 0.08 - 0.513; IGK = 0.244, 95%-CI = 0.058 - 0.466; IGL = 0.197, 95%-CI = 0.014 - 0.401; TRA = 0.126, 95%-CI = 0.017 - 0.241; TRB = 0.18, 95%-CI = 0.036 - 0.329). (**B**) Intraindividual comparison of Simpson clonality across LNs and pathologic assessment. Red lines are the median of individual medians. (**C**) Volcano plot showing differentially expressed genes in melanoma-positive LNs ( *PMEL* logFC = 2.088, q = 7.316e-03; *MLANA* logFC = 3.147, q = 9.664e-03; *TYR* logFC = 3.538, q = 3.056e-03; *DCT* logFC = 2.842, q = 8.342e-03; *SOX10* logFC = 2.76, q = 8.258e-03; *PRAME* logFC = 1.64, q = 2.724e-02; *HIST1H1A* logFC = 1.299, q = 9.265e-04; *IFITM2* logFC = −0.582, q = 2.982e-03; *CISH* logFC = −0.612, q = 2.982e-03; *JAK1* logFC = −0.489, q = 8.342e-03; *JAK2* logFC = −0.588, q = 9.105e-03). (**D**) Upregulated GO terms in melanoma-positive LNs. (**E**) Downregulated GO terms in melanoma-positive LNs. Nodes are colored by p value and labelled with the number of differentially expressed genes per GO term.

**Figure 2:**
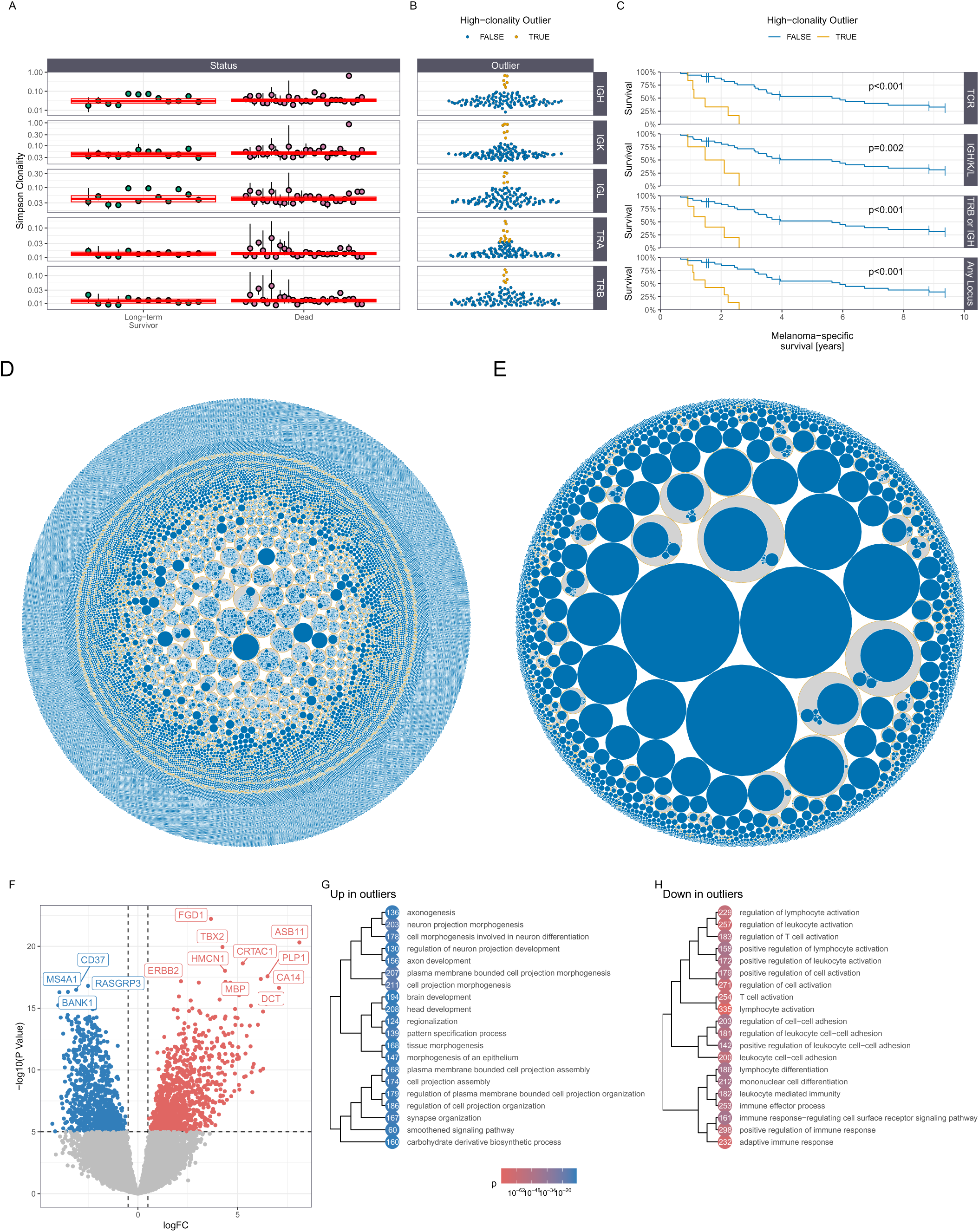
Patients with LNs with very high immune repertoire clonality have a worse prognosis. (**A**) Comparison of Simpson clonality by follow-up status from the melanoma registry. Circles indicate the patient median and whiskers the intraindividual range. Red lines show the model’s conditional effects with credibility intervals (IGH = 0.083, 95%-CI = −0.225 - 0.381; IGK = 0.118, 95%-CI = −0.126 - 0.363; IGL = 0.009, 95%-CI = −0.278 - 0.284; TRA = 0.023, 95%-CI = −0.142 - 0.196; TRB = 0.036, 95%-CI = −0.158 - 0.24). (**B**) Classification of high-clonality outliers based on a log_10_(clonality) that is > 3x the median absolute deviation above the median. (**C**) Melanoma-specific survival. Patients were classified as high-clonality outliers if at least one LN had a high-clonality outlier in the referenced loci. P values were determined using a log-rank test. (**D** & **E**) Exemplary TRB repertoire representation of lymph nodes 381 (negative, no outlier) and 380 (positive, outlier), both of patient hist11. Blue circles depict unique sequences, scaled by their clone size. Yellow circles group together clusters, as defined by clusTCR. (**F**) Volcano plot showing differentially expressed genes in outlier versus non-outlier LNs. *MS4A1* [CD20] logFC = −3.946, q = 5.231e-14; *CD19* logFC = −3.334, q = 3.710e-12; *CD79A* logFC = −3.379, q = 3.100e-06; *CD3D* logFC = −3.202, q = 7.853e-10; *CD4* logFC = −1.596, q = 5.966e-06; *CD8A* logFC = −1.37, q = 6.017e-04; *CD40LG* logFC = −3.469, q = 2.293e-10; *CD69* logFC = −3.09, q = 2.382e-10; *DCT* logFC = 6.792, q = 9.400e-14; *PMEL* logFC = 5.096, q = 1.061e-07; *MLANA* logFC = 6.289, q = 3.956e-09; *PRAME* logFC = 5.067, q = 6.224e-10; *SOX10* logFC = 5.746, q = 1.598e-07; *BCAN* logFC = 6.445, q = 3.302e-13; *MITF* logFC = 3.979, q = 3.775e-11; *ERBB3* logFC = 5.33, q = 1.418e-08. (**G**) Upregulated GO terms in outlier LNs. (**H**) Downregulated GO terms in outlier LNs.

The expression of MDAs and melanoma markers was higher in melanoma-positive LNs (**Figure 1C**). Metastatic LNs also showed upregulation of cell cycle associated transcripts such as *HIST1H1A* (log2 fold change (logFC) = 1.299, q = 9.265e-04, **Figure 1C**). In contrast, interferon signaling (*IFITM2* logFC = −0.582, q = 2.982e-03), NK cell activity (*CISH* logFC = −0.612, q = 2.982e-03) and immune system activation via Janus kinase signaling were downregulated (*JAK1* logFC = −0.489, q = 8.342e-03; *JAK2* logFC = −0.588, q = 9.105e-03 **Figure 1C**). This pattern is further reflected by the upregulation of gene ontology (GO) terms linked to pigmentation, neurogenesis (reflecting the neuroectodermal origin of melanocytes) and cell division and downregulation of immune system activation-related GO terms (**Figure 1D-E**).

Clonality showed no consistent association with tumor thickness, nodal stage or overall UICC stage across loci (**Supplemental Figure 1**).

### High-clonality outliers are associated with poor patient outcome

Assuming that LNs reflect the status of the locoregional immune system, we hypothesized that changes in immune receptor clonality might be associated with survival. Overall, clonality did not differ between patients who died and long-term survivors (IGH = 0.083, 95%-CI = −0.225 - 0.381; IGK = 0.118, 95%-CI = −0.126 - 0.363; IGL = 0.009, 95%-CI = −0.278 - 0.284; TRA = 0.023, 95%-CI = −0.142 - 0.196; TRB = 0.036, 95%-CI = −0.158 - 0.24; **Figure 2A**). However, we observed that high-clonality outliers were more frequent among patients who later died (**Figure 2A**). Therefore, we defined LNs with clonality values greater than three times the median absolute deviation (MAD) above the median overall clonality as outliers (**Figure 2B** and **Supplemental Figure 2**). High-clonality outliers in either a TCR locus (p<0.001), a BCR locus (p=0.002), the TRB or IGH locus (p<0.001) or across any locus (p<0.001) were associated with significantly poorer melanoma-specific survival (**Figure 2C**). Outliers showed a marked reduction in B and T cell markers (*MS4A1* logFC = −3.946, q = 5.231e-14; *CD19* logFC = −3.334, q = 3.710e-12; *CD79A* logFC = −3.379, q = 3.100e-06; *CD3D* logFC = −3.202, q = 7.853e-10; *CD4* logFC = −1.596, q = 5.966e-06; *CD8A* logFC = −1.37, q = 6.017e-04), as well as activation markers (*CD40LG* logFC = −3.469, q = 2.293e-10; *CD69* logFC = −3.09, q = 2.382e-10, **Figure 2F**). This can visually be exemplified by LNs 380 and 381 of patient hist11, where the outlier LNs shows a prominent restriction of the TRB repertoire (**Figure 2D-E**). MDAs and melanoma-associated transcription factors on the other hand were high (*DCT* logFC = 6.792, q = 9.400e-14; *PMEL* logFC = 5.096, q = 1.061e-07; *MLANA* logFC = 6.289, q = 3.956e-09; *PRAME* logFC = 5.067, q = 6.224e-10; *SOX10* logFC = 5.746, q = 1.598e-07; *BCAN* logFC = 6.445, q = 3.302e-13; *MITF* logFC = 3.979, q = 3.775e-11; *ERBB3* logFC = 5.33, q = 1.418e-08 **Figure 2F**). Consistently, gene ontology analysis demonstrated enrichment of neurogenesis-related terms and the downregulation of immune system-related terms in high-clonality outliers (**Figure 2F-H**).

### Validation cohort

To validate these findings, we recruited a second, contemporary validation cohort of 87 melanoma patients undergoing sentinel LN biopsy between 2022 and 2024 (characteristics in **Supplementary Table 2**). We sequenced 70 LNs and 54 matched peripheral blood mononuclear cell (PBMC) samples. Melanoma-positive LNs showed an increased Simpson clonality for IGH (0.406, 95%-CI = 0.07 - 0.721), IGK (0.384, 95%-CI = 0.075 - 0.676) and TRB (0.291, 95%-CI = 0.037 - 0.612), whereas for IGL (0.237, 95%-CI = −0.055 - 0.523) and TRA (0.193, 95%-CI = −0.014 - 0.472), credibility intervals included zero (**Figure 3A**). No difference could be observed between the immune repertoire clonality of PBMCs of patients with versus without lymph node involvement (IGH = 0.136, 95%-CI = −0.17 - 0.461; IGK = −0.012, 95%-CI = −0.23 - 0.212; IGL = −0.01, 95%-CI = −0.212 - 0.208; TRA = 0.102, 95%-CI = −0.329 - 0.541; TRB = 0.253, 95%-CI = −0.241 - 0.757; **Figure 3B**).

**Figure 3:**
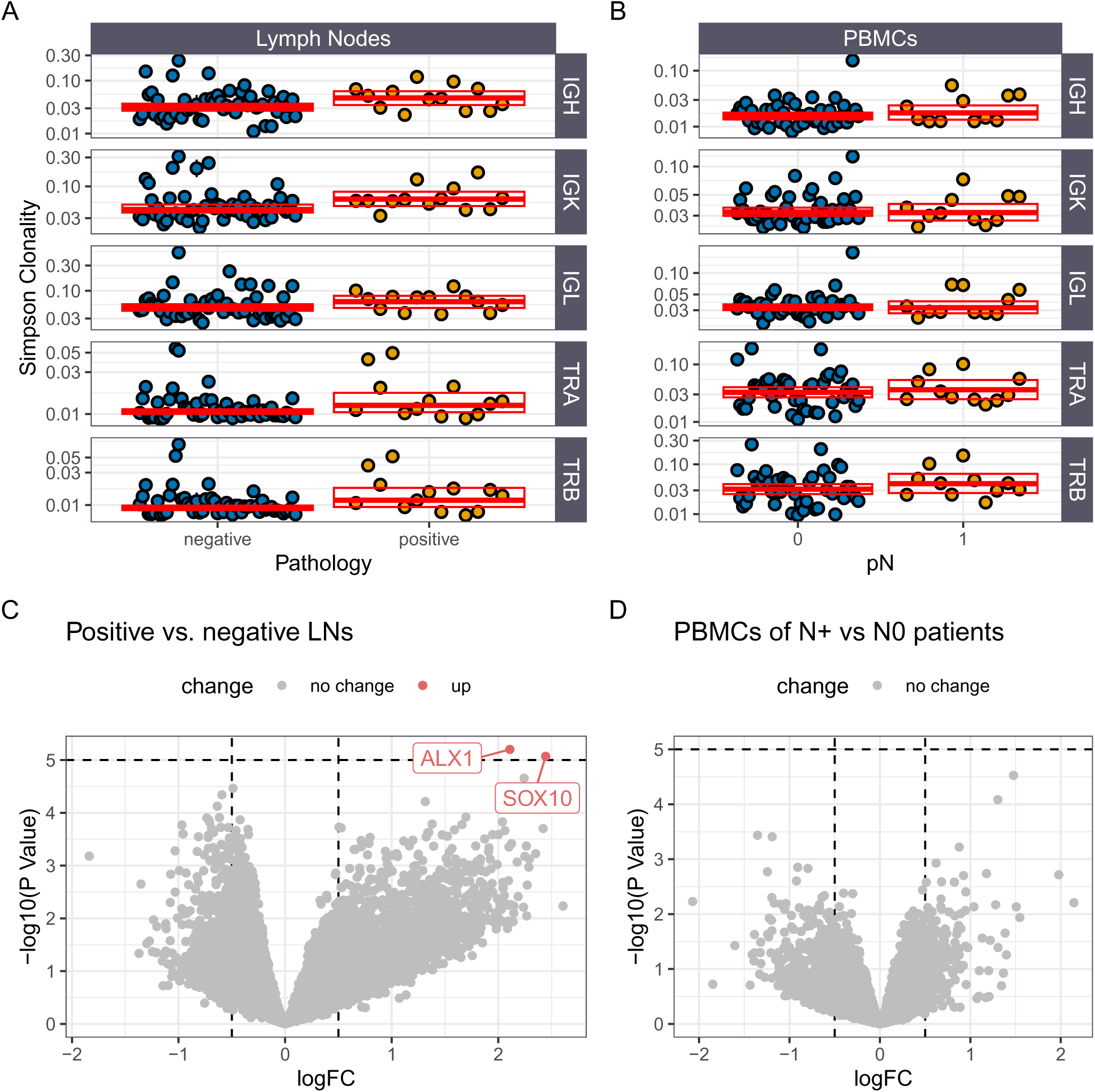
Effect of melanoma spread on lymph node immune repertoire clonality and gene expression in the validation cohort. (**A**) Comparison of Simpson clonality in melanoma-positive versus -negative LNs for IGH, IGK and TRB but not IGL and TRA ( IGH = 0.406, 95%-CI = 0.07 - 0.721; IGK = 0.384, 95%-CI = 0.075 - 0.676; TRB = 0.291, 95%-CI = 0.037 - 0.612; IGL = 0.237, 95%-CI = −0.055 - 0.523; TRA = 0.193, 95%-CI = −0.014 - 0.472). (**B**) Contrasts clonality in PBMCs of patients with versus without lymphonodal spread ( IGH = 0.136, 95%-CI = −0.17 - 0.461; IGK = −0.012, 95%-CI = −0.23 - 0.212; IGL = −0.01, 95%- CI = −0.212 - 0.208; TRA = 0.102, 95%-CI = −0.329 - 0.541; TRB = 0.253, 95%-CI = −0.241 - 0.757). (**C**) Volcano plot showing differentially expressed genes in melanoma-positive versus -negative LNs. *SOX10* logFC = 2.444, q = 5.608e-02; *ALX1* logFC = 2.108, q = 5.608e-02. (**D**) Volcano plot showing differentially expressed genes in PBMCs of patients with versus without lymphonodal spread. In A & B circles indicate the patient median and whiskers the intraindividual range and red lines show the model’s conditional effects with credibility intervals.

At the gene expression level, only two genes showed a trend toward upregulation in positive versus negative LNs, although neither reached the false discovery rate (FDR)<0.05 significance threshold (*SOX10* logFC = 2.444, q = 5.608e-02; *ALX1* logFC = 2.108, q = 5.608e-02; **Figure 3C**), whereas no difference was found in the gene expression profiles of N0 vs N1 patients (**Figure 3D**).

### Melanocyte differentiation antigen-directed T cell responses in peripheral blood and LNs

To assess T cell reactivity against well-known MDAs, we performed T cell stimulation assays using protein-spanning peptide pools (PSPs) against gp100, melan-A, tyrosinase, tyrosinase-related protein 2 (TYRP2), and a pool of known immunogenic peptides from CMV, EBV and influenza (CEF) as positive control.

A total of 121 patients were included, 30 (24.8%) of whom had detectable melanoma cell spread to the LN. In 101 cases, single LNs were acquired, whereas 15 and 3 patients had 2 and 3 LNs available, respectively. 140 LNs of 119 patients were stimulated. **Figure 4** shows the stimulation results separately for CD4+ and CD8+ cells for each peptide pool. Stimulation results are displayed in a circular fashion, where each patient is one column and the available materials (PBMCs and LNs) are rows of the circles. The sectors are split so that patients are grouped together as follows: (i) patients who have shared responses in PBMCs and LNs, (ii) those with exclusive activation in one compartment and (iii) those without activation.

**Figure 4:**
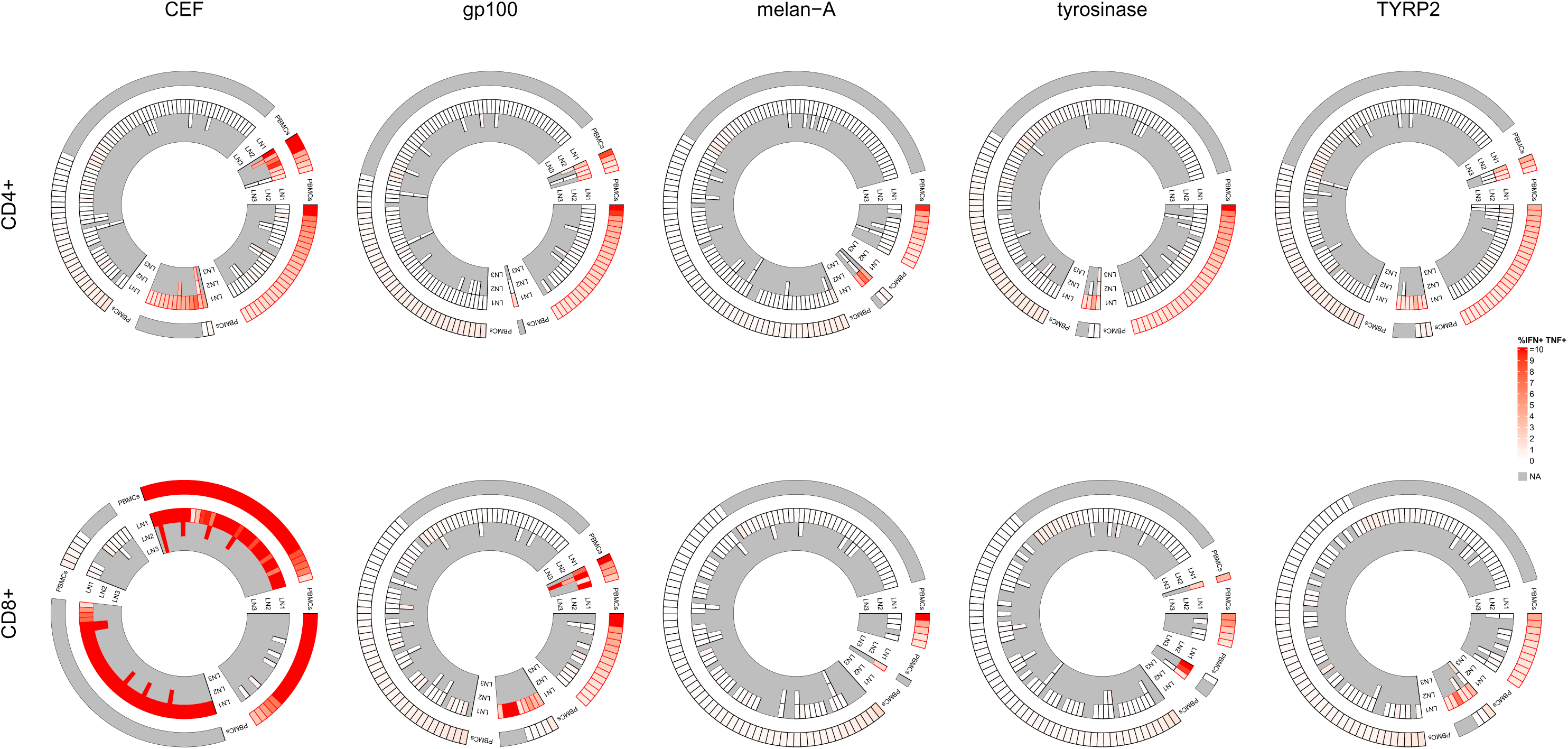
T cell stimulation of patient LNs and PBMCs with MDA PSPs. For each PSP, CD4 and CD8 responses are shown in a circular layout where the columns are patients and the rows are the available materials, colored by the interferon γ and TNF positivity following 10-day T cell expansion and stimulation with PSPs. CEF was used as positive control. Red cell borders indicate activation as defined by a frequency of TNF+ IFN-γ+ cells of > 1% and >2x the medium control.

PBMCs were stimulated from 69 patients, 67 of whom had matched LNs and PBMCs available. In general, PBMCs were more frequently activated than LNs across stimulation conditions and T cell subsets but the differences were more pronounced in CD4+ than in CD8+ T cells (**Supplemental Figure 3**). MDA-directed reactivity was detected in 19 (13.6%, CD4+) and 23 (16.4%, CD8+) LNs. CD4+ responses were directed against TYRP2 (10 LN), gp100 (5 LN), tyrosinase (5 LN) and melan-A (3 LN), whereas CD8+ responses targeted gp100 (15 LN), TYRP2 (6 LN), tyrosinase (4 LN) and melan-A (1 LN). The rate was higher in PBMCs with 52 (75.4%, CD4+) and 31 (44.9%, CD8+) patients responding. Here, responses were directed against TYRP2 (29 patients), tyrosinase (29 patients), gp100 (28 patients) and melan-A (11 patients) in CD4+ cells and gp100 (19 patients), TYRP2 (11 patients), tyrosinase (6 patients) and melan-A (5 patients) in CD8+ cells. When reactivity was recorded, it was shared against the same peptide pool between PBMCs and LNs in 4 cases for both CD4+ and CD8+ cells stimulated with a gp100 PSP, 3 cases for CD4+ cells stimulated with TYRP2 and 1 case for CD4+ cells stimulated with a tyrosinase PSP (**Figure 4**). Reactivity was concordant between CD4+ and CD8+ cells in 2 and 15 of activated samples. Concordant activation in multiple LNs of the same patient was observed once for CD8+ cells in gp100 stimulated LNs and once for CD4+ cells in tyrosinase stimulated LNs. Intriguingly, patient val101, who had a shared CD8+ T cell response against gp100 across compartments, showed marked regression zones in their primary tumor.

## Discussion

In this study, we leveraged two large cohorts comprising a total of 273 lymph nodes across 195 patients with melanoma — one from the pre-ICI era and one a contemporary validation cohort — to investigate how immune repertoire clonality in LNs correlates with disease progression and patient survival and how MDA-directed immunity is shared between tissues.

Our findings revealed that increased TCR and BCR clonality in LNs, especially when manifesting as high-clonality outliers, is associated with reduced immune activity and poor clinical outcomes. We also found that melanoma-positive LNs exhibit higher clonality than tumor-free nodes. Notably, increased expression of MDAs and concurrent downregulation of interferon and immune activation pathways in these nodes suggest that the elevated clonality may not reflect effective anti-tumor immunity, but rather immunosuppression or exhaustion and metastatic outgrowth. This is in line with a prior report of increased TRB clonality alongside a phenotypically more immunotolerant environment in melanoma-positive versus -negative sentinel LNs (25) and also confirms prior observations that a higher TRB clonality is associated with worse patient outcomes, if no ICI is administered (26). It further accords with the recent finding that a tolerogenic program in tumor-draining LNs—medullary sinus macrophage–derived IL-33 activating regulatory T cells—impairs anti-tumor immunity and, in melanoma, scales with nodal stage and survival (10), consistent with our interpretation that the high-clonality, immune-depleted outlier nodes represent a locally suppressed rather than a productively reactive compartment. Interestingly, the increase in clonality was not uniform across all patients, indicating interindividual variability in immune responsiveness or tumor influence. In fact, the clonality increase and transcriptional changes in positive samples were the smallest in the contemporary validation cohort (where all but one metastasis was detected by quantitative immunocytology (27, 28) and missed in conventional routine pathology), more apparent in the full pre–ICI era cohort and most pronounced when comparing high-clonality outliers with non-outliers, a comparison where also transcriptional data showed a steep increase in melanocytic transcripts even when correcting the differential expression model for lymph node positivity. Therefore, the increase in clonality is likely associated with tumor outgrowth, corresponding to the escape stage of the three Es model of cancer immunoediting (29). These observations align with the hypothesis that effective tumor immunity relies on a diverse and polyclonal T cell population capable of recognizing a wide range of antigens (30). In contrast, highly clonal repertoires may represent exhausted or tumor-specific populations that have failed to control tumor progression (31). Such skewed repertoires may also reflect tumor-induced expansion of ineffective clones or the collapse of broader immune surveillance.

To map MDA-directed immunity, we stimulated the PBMCs and sentinel LN-derived T cells of 121 patients with PSPs of four highly immunogenic MDAs. All four tested MDAs elicited responses, with TYRP2 and gp100 being most frequently recognized. Intriguingly, the immune responses were infrequently shared across lymph nodes of the same individual or across compartments. Similarly, Oliveira et al. found MDA-reactive tumor-infiltrating lymphocytes (TILs) that were in an exhausted transcriptional state and infrequently shared with the patient’s peripheral blood T cell pool (32). Together, these results suggest a strong compartmentalization of the immune response and underscore the importance of tissue-resident memory T cells (5). Notably, the single patient with a CD8+ gp100 response shared between peripheral blood and LN (val101) presented with marked regression zones in the primary tumor, anecdotally linking cross-compartment MDA-directed immunity to clinically evident anti-tumor activity.

These findings are highly relevant in the context of current debates regarding the surgical management of tdLNs in melanoma (16). While complete LN dissection does not improve survival in sentinel LN-positive patients (12), the immunological importance of tdLNs is increasingly recognized in the era of ICI therapy (13, 14). Our results suggest that the immunological state of tdLNs, particularly repertoire clonality, might serve as a biomarker for prognosis and treatment stratification.

Further, since our pre–ICI era cohort predates the widespread use of ICI therapy, it provides a unique window into the natural history of immune-tumor interactions in melanoma. As ICI therapies become standard, it will be important to determine whether pre-treatment LN clonality or the emergence of high-clonality outliers can predict therapeutic responses or resistance mechanisms.

Our study is limited by its retrospective nature and by the single-center origin of both cohorts, which may restrict generalizability.

Additionally, while Simpson clonality is a robust and interpretable metric, it cannot distinguish between effective and ineffective clonal expansions. Future studies incorporating functional assays, single-cell analyses, or antigen specificity profiling could provide more mechanistic insight into the observed clonality patterns.

## Methods

### Sex as a biological variable

Our study examined samples from both male and female patients (see Supplementary Table 1 and Supplementary Table 2). Sex was not used as a selection criterion, and both sexes were included in all analyses. Our findings are expected to be relevant for both sexes; the cohorts were not powered to detect sex-specific differences.

### Human samples

#### Pre–immune checkpoint inhibitor era cohort

All LNs were obtained between 1994 and 2002 from sentinel LN biopsies and elective lymphadenectomies in a previous study assessing the use of quantitative polymerase chain reaction (PCR) for the detection of melanoma LN metastasis (33). LN fragments were stored snap frozen at −80°C.

#### Validation cohort

All LNs were obtained from sentinel LN biopsy as leftover of the routinely performed quantitative immuno-cytology (27) between 2022 and 2024. PBMCs were collected prospectively from participating patients at the time of sentinel LN biopsy.

### Sequencing

LNs were homogenized in 600µl buffer RA1 (MACHEREY-NAGEL, Cat #: 740955.250) and 6µl β-mercaptoethanol (CARL ROTH, Cat #: 4227.1) using a scalpel (pfm medical, Cat #: 200130010) and pestle (BelArt, Cat #: F65000-0006) and incubated at room temperature for 15 minutes. Thereafter, the lysate was filled to 1.5ml with RA1 and processed following the manufacturer’s recommendations using the NucleoSpin RNA Mini Kit (MACHEREY-NAGEL, Cat #: 740955.250). Resulting RNA was eluted in 60µl of nuclease-free water and quantified on an IMPLEN NanoPhotometer® NP80. TCR and BCR libraries were generated using the DriverMap™ AIR TCR/BCR Profiling Kit (human RNA, Cellecta, Cat #: DMAIR-HTBR-96). Gene expression sequencing libraries were generated using the DriverMap™ Human Genome-Wide Expression Profiling Kit, V2 (Cellecta, Cat #: DM2-HGW-96).

All libraries were subjected to quality control and sequencing at CeGaT (Tübingen, Germany) at a depth of 6-10 million reads per sample.

### Sequencing data analysis

#### TCR/BCR sequencing

Raw FASTQ sequence files generated from TCR and BCR libraries were processed using the mixcr analyze cellecta-human-rna-xcr-umi-drivermap-air preset in MiXCR version 4.7 (34).

We used Simpson clonality as a measure of clonality with the definition Simpson clonality = 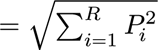, where *R* is the number of unique rearrangements and *P_i_* is the relative fraction of the i^th^ rearrangement.

To account for differences in sequencing depth, samples were downsampled to 0.5 ∗ D2 unique mole-cular identifiers (UMIs), where D2 is the second decile. Downsampling was performed as 1000 random bootstraps with replacement and calculation of Simpson clonality was performed on all, with the mean value being used in further analyses.

TRB sequences were clustered by clusTCR (v. 1.0.3+3.g5fa6b46) (35).

#### Gene expression libraries

A custom decoy-aware salmon (36) index was built using the amplicon sequences provided by Cellecta and the Homo_sapiens.GRCh38.112 reference with k-mers of length 19. Raw FASTQ sequence files were quantified in mapping-based mode in salmon (v. 1.10.3) (36) using `salmon quant` with the custom reference. Enriched GO terms were identified using limma’s `goana` function with a FDR threshold of 0.01 and filtered to BP terms of sizes smaller than 800. They were clustered using hierarchical clustering of Wang similarity measures as implemented in GOSemSim (v. 2.36.0) (37).

### T cell stimulation assay

#### PBMC separation and storage

PBMCs were separated from 50ml of EDTA blood using Pancoll human (PAN Biotech, Cat#: P04-601000) and either used directly for T cell culture or frozen in Cryostor CS10 (Stemcell Cat #: 7930), according to the manufacturer’s instructions.

#### LN cell storage

Surplus cells available from routine quantitative immunocytology (27) were frozen in Cryostor CS10 (Stemcell Cat #: 7930).

#### T cell culture

Cells were stimulated with custom-made MDAs PSPs of 15mers with an overlap of 11AA or a CEF pool (all by GenScript), at a concentration of 2µg/ml/peptide. They were subsequently expanded with IL-2 (Novartis), restimulated with peptide pools on day 10 with added brefeldin A (Sigma-Aldrich, Cat #: B7651). After 6 hours of incubation, samples were stained with BioLegend antibodies for dead cells (Zombie Aqua, Cat #: 423101), CD3 (Cat #: 317341), CD4 (Cat #: 317413), CD8 (Cat #: 344707), CD45RA (Cat #: 304129), IFN-γ (Cat #: 506506), TNF (Cat #: 502913) and measured on a Cytek Aurora spectral cytometer. Samples were classified as reactive when the frequency of TNF+ IFN-γ+ cells was > 1% of the parent CD4 or CD8 gate and >2x the medium control. Samples with CD4 or CD8 counts less than 500 were excluded from analysis.

### Statistics

Unless otherwise specified, modeling was performed using the brms package (v. 2.23.0) in R (v. 4.5.2) with model specifications as detailed in the supplementary information (38). Coefficients are reported with their posterior mean and 95% credibility interval (95%-CI). Survival analysis was performed using the Kaplan–Meier estimator and groups were compared using the log-rank test. High-clonality outliers were defined per immune receptor locus as LNs whose log-transformed Simpson clonality exceeded the locus median by more than three times the MAD; patients were classified as outliers if at least one of their LNs met this criterion in the referenced loci. Differential gene expression testing was performed using limma’s (v. 3.66.0) `voomLmFit` and `eBayes` functions; genes with an FDR below 0.05 were considered significant. Plots were generated using ggplot2 (v. 4.0.1) (39) and circlize (v. 0.4.17) (40).

### Study approval

The use of archival tissue from the pre–ICI era cohort was approved by the ethics commission of the University of Tübingen (589/2022BO2); patients had consented to the use of their tissue for research purposes. The collection and analysis of samples from the contemporary validation cohort was approved by the ethics commission of the University of Tübingen (118/2021BO2) and all patients provided written informed consent prior to inclusion.

### Data availability

Sequencing data and patient-level metadata are available from Zenodo at doi.org/10.5281/zen-odo.18776328. Values for all data points shown in graphs and reported in summary statistics are provided in the Supporting Data Values file.

## Author contributions

Conceptualization: VW, LF, Methodology: VW, TS, LF, Formal Analysis: VW, Investigation: VW, Resources: AU, CS, LF, Data Curation: VW, JH, SAF, ST, MV, MK, LN, ULS, Writing – Original Draft: VW, LF, Writing – Review & Editing: All authors, Visualization: VW, Supervision: LF, CS, Project Administration: LF, Funding Acquisition: LF

## Acknowledgments

The authors thank the patients who consented to the use of their tissue and clinical data for this study and the team of the Department of Dermatology biobank at the University Hospital Tübingen for sample curation. This work was supported by the de.NBI Cloud within the German Network for Bioinformatics Infrastructure (de.NBI) and ELIXIR-DE (Forschungszentrum Jülich and W-de.NBI-001, W-de.NBI-004, W-de.NBI-008, W-de.NBI-010, W-de.NBI-013, W-de.NBI-014, W-de.NBI-016, W-de.NBI-022).

## Abbreviations

95%-CI: 95% credibility interval
TYRP2: tyrosinase-related protein 2
BCR: B cell receptor
CDR3: complementarity determining region 3
CEF: CMV, EBV and influenza
FDR: false discovery rate
GO: gene ontology
ICI: immune checkpoint inhibitor
IG: immunoglobulin
IGH: IG heavy chain
IGK: IG κ chain
IGL: IG λ chain
logFC: log2 fold change
LN: lymph node
MAD: median absolute deviation
MDA: melanocyte differentiation-antigen
PBMC: peripheral blood mononuclear cell
PCR: polymerase chain reaction
PSP: protein-spanning peptide pool
TCR: T cell receptor
tdLN: tumor-draining lymph node
TIL: tumor-infiltrating lymphocyte
TRA: TCR α chain
TRB: TCR β chain
UMI: unique molecular identifier

## Figures

**Supplemental Figure 1:**
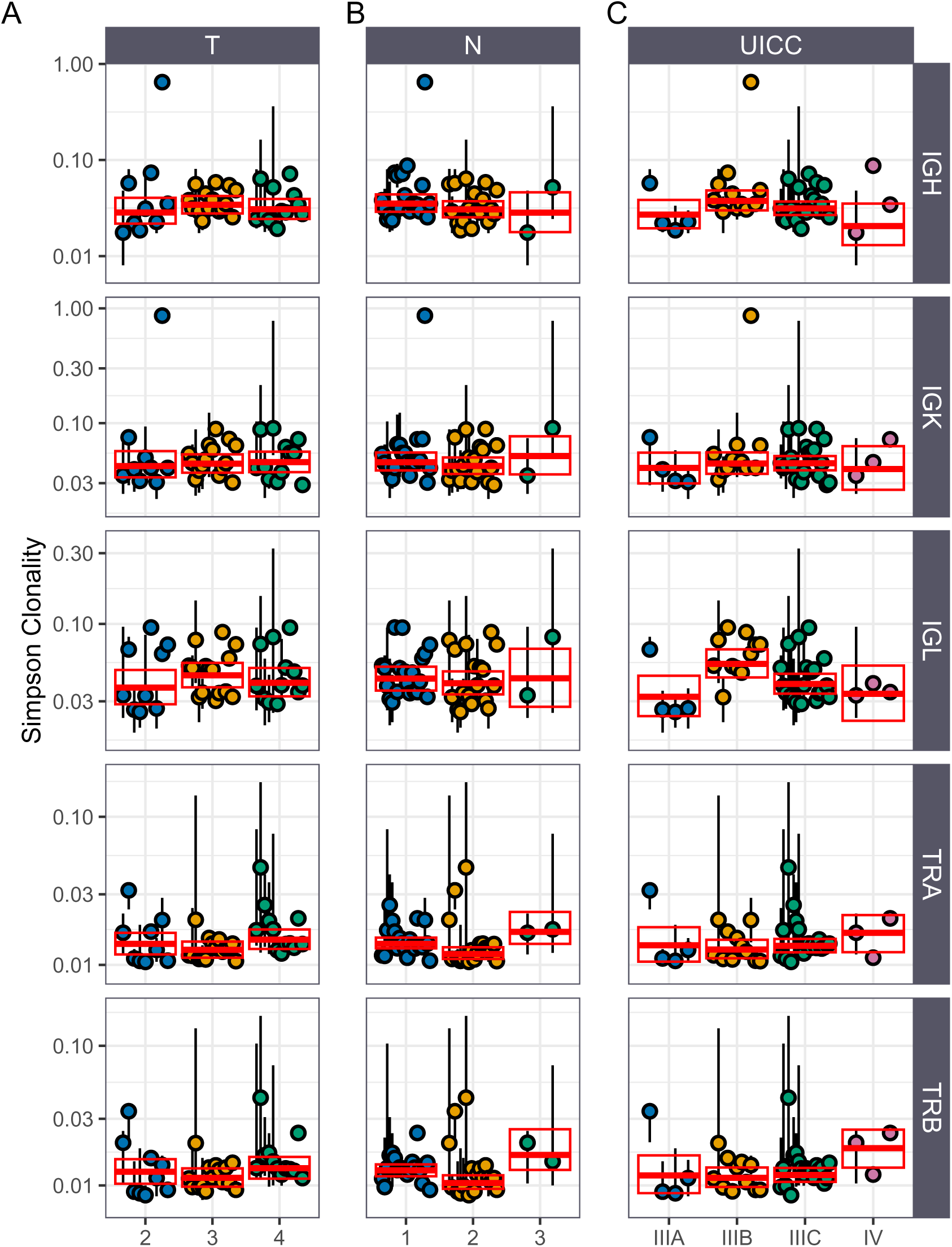
Comparison of Simpson clonalities across metrics of tumor spread. (**A**) compares clonality across different primary tumor thicknesses. (**B**) contrasts LN clonality according to overall LN spread. (**C**) shows LN clonality according to overall UICC stage. Red lines are the median of individual medians. Circles indicate the patient median and whiskers the intraindividual range.

**Supplemental Figure 2:**
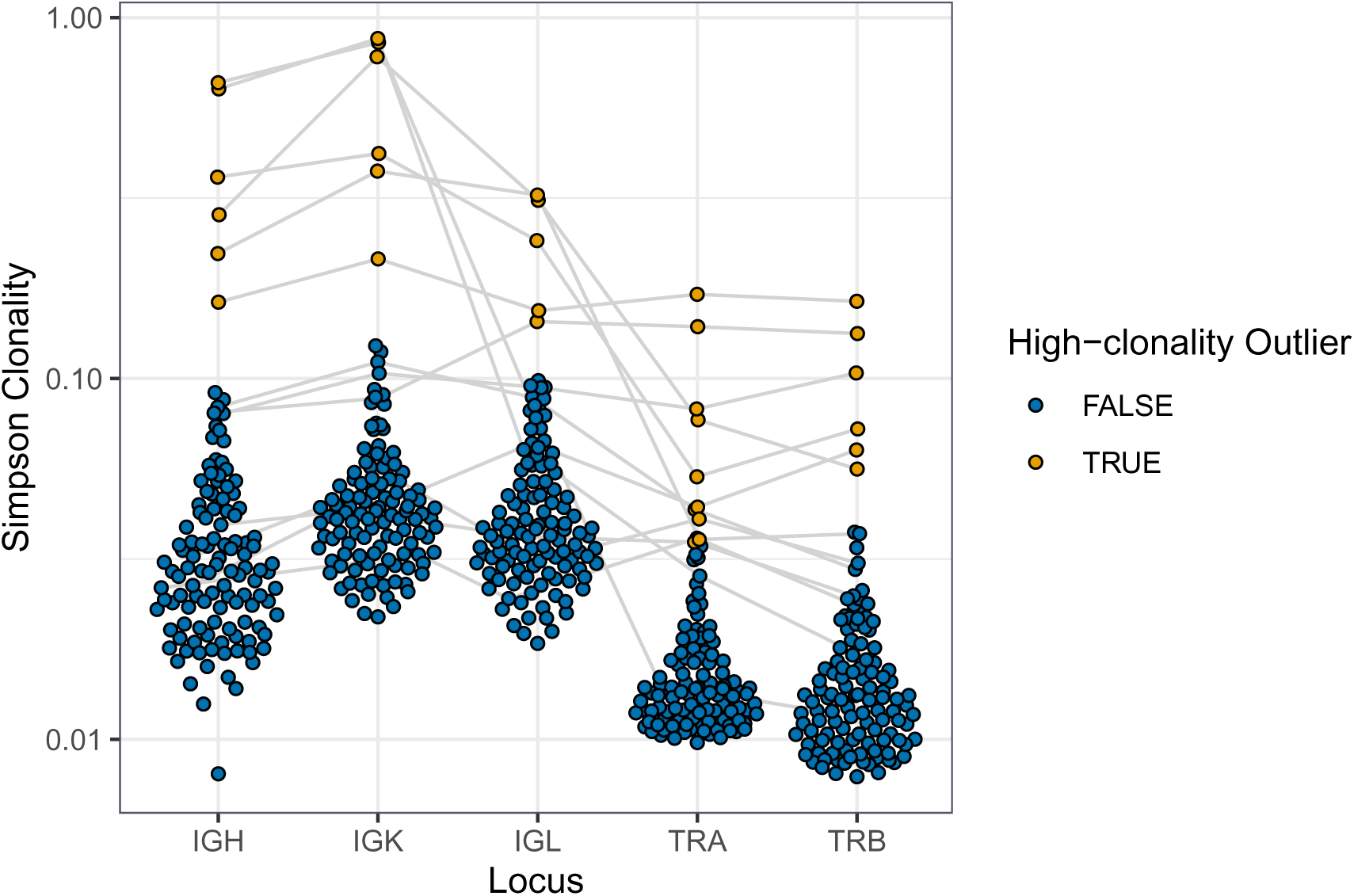
High-clonality outliers connected by LN sample.

**Supplemental Figure 3:**
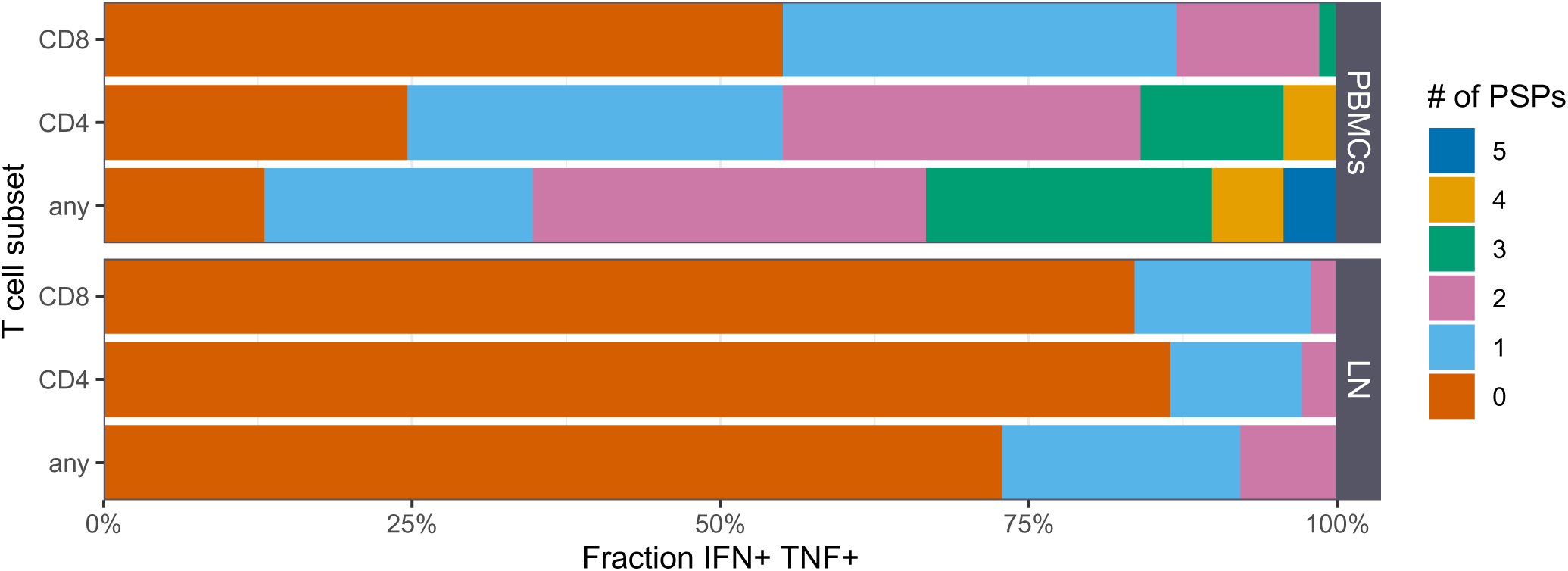
Comparison of the fraction of activated samples by T cell subsets colored by the number of PSPs eliciting this reaction.

## Supplement

**Supplemental Table 1:**
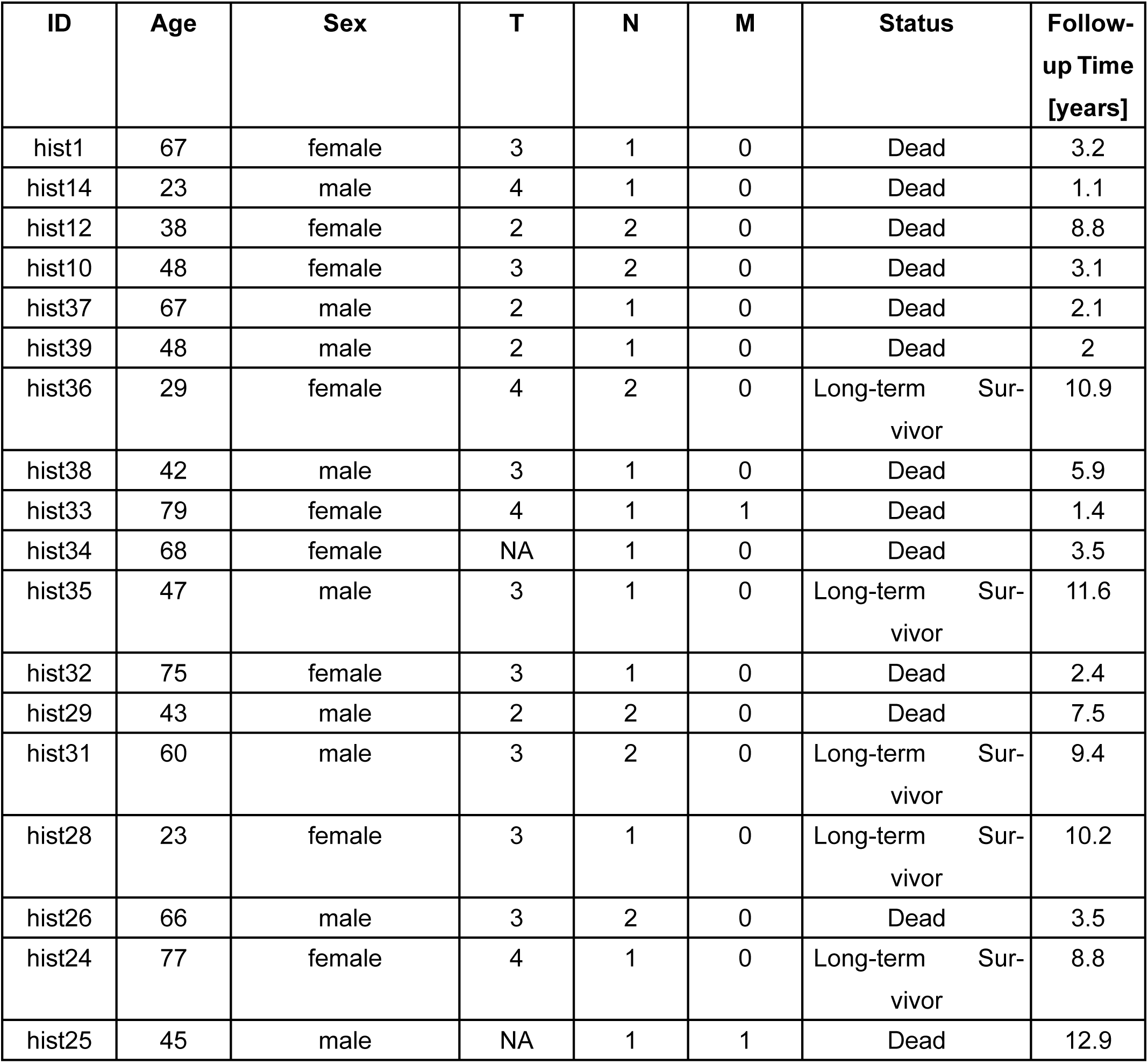
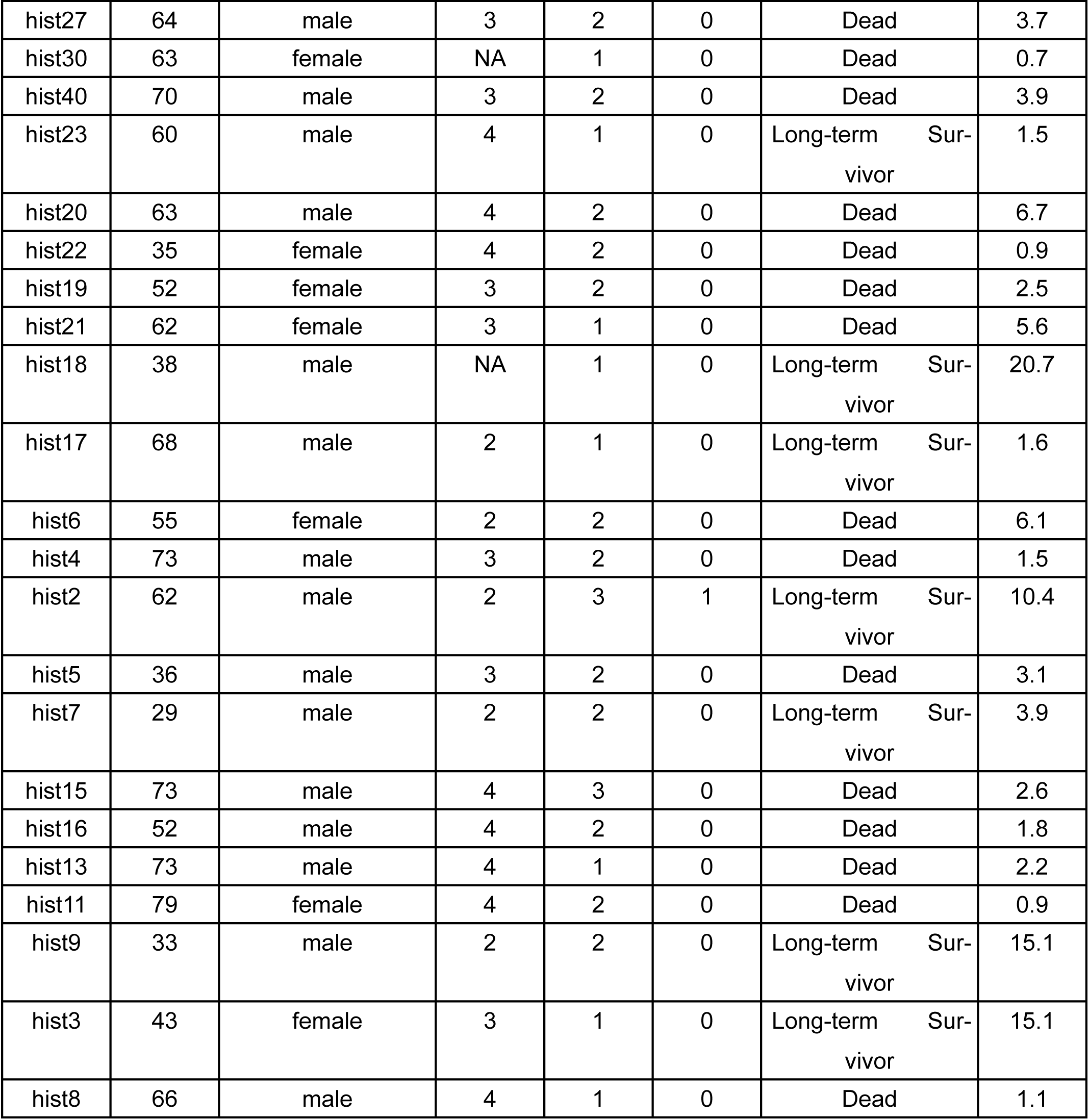
pre–ICI patient cohort characteristics.

| ID | Age | Sex | T | N | M | Status | Follow-up Time [years] |
| --- | --- | --- | --- | --- | --- | --- | --- |
| hist1 | 67 | female | 3 | 1 | 0 | Dead | 3.2 |
| hist14 | 23 | male | 4 | 1 | 0 | Dead | 1.1 |
| hist12 | 38 | female | 2 | 2 | 0 | Dead | 8.8 |
| hist10 | 48 | female | 3 | 2 | 0 | Dead | 3.1 |
| hist37 | 67 | male | 2 | 1 | 0 | Dead | 2.1 |
| hist39 | 48 | male | 2 | 1 | 0 | Dead | 2 |
| hist36 | 29 | female | 4 | 2 | 0 | Long-term Survivor | 10.9 |
| hist38 | 42 | male | 3 | 1 | 0 | Dead | 5.9 |
| hist33 | 79 | female | 4 | 1 | 1 | Dead | 1.4 |
| hist34 | 68 | female | NA | 1 | 0 | Dead | 3.5 |
| hist35 | 47 | male | 3 | 1 | 0 | Long-term Survivor | 11.6 |
| hist32 | 75 | female | 3 | 1 | 0 | Dead | 2.4 |
| hist29 | 43 | male | 2 | 2 | 0 | Dead | 7.5 |
| hist31 | 60 | male | 3 | 2 | 0 | Long-term Survivor | 9.4 |
| hist28 | 23 | female | 3 | 1 | 0 | Long-term Survivor | 10.2 |
| hist26 | 66 | male | 3 | 2 | 0 | Dead | 3.5 |
| hist24 | 77 | female | 4 | 1 | 0 | Long-term Survivor | 8.8 |
| hist25 | 45 | male | NA | 1 | 1 | Dead | 12.9 |
| hist27 | 64 | male | 3 | 2 | 0 | Dead | 3.7 |
| hist30 | 63 | female | NA | 1 | 0 | Dead | 0.7 |
| hist40 | 70 | male | 3 | 2 | 0 | Dead | 3.9 |
| hist23 | 60 | male | 4 | 1 | 0 | Long-term Survivor | 1.5 |
| hist20 | 63 | male | 4 | 2 | 0 | Dead | 6.7 |
| hist22 | 35 | female | 4 | 2 | 0 | Dead | 0.9 |
| hist19 | 52 | female | 3 | 2 | 0 | Dead | 2.5 |
| hist21 | 62 | female | 3 | 1 | 0 | Dead | 5.6 |
| hist18 | 38 | male | NA | 1 | 0 | Long-term Survivor | 20.7 |
| hist17 | 68 | male | 2 | 1 | 0 | Long-term Survivor | 1.6 |
| hist6 | 55 | female | 2 | 2 | 0 | Dead | 6.1 |
| hist4 | 73 | male | 3 | 2 | 0 | Dead | 1.5 |
| hist2 | 62 | male | 2 | 3 | 1 | Long-term Survivor | 10.4 |
| hist5 | 36 | male | 3 | 2 | 0 | Dead | 3.1 |
| hist7 | 29 | male | 2 | 2 | 0 | Long-term Survivor | 3.9 |
| hist15 | 73 | male | 4 | 3 | 0 | Dead | 2.6 |
| hist16 | 52 | male | 4 | 2 | 0 | Dead | 1.8 |
| hist13 | 73 | male | 4 | 1 | 0 | Dead | 2.2 |
| hist11 | 79 | female | 4 | 2 | 0 | Dead | 0.9 |
| hist9 | 33 | male | 2 | 2 | 0 | Long-term Survivor | 15.1 |
| hist3 | 43 | female | 3 | 1 | 0 | Long-term Survivor | 15.1 |
| hist8 | 66 | male | 4 | 1 | 0 | Dead | 1.1 |

**Supplemental Table 2:** Contemporary validation cohort patient characteristics.

| id | age | sex | pT | pN | cM |
| --- | --- | --- | --- | --- | --- |
| val123 | 48 | male | 2 | 0 | 0 |
| val131 | 70 | male | 2 | 0 | 0 |
| val54 | 51 | female | 4 | 1 | 0 |
| val61 | 76 | male | 2 | 0 | 0 |
| val26 | 70 | female | 1 | 0 | 0 |
| val101 | 52 | male | 1 | 0 | 0 |
| val102 | 73 | female | 2 | 0 | 0 |
| val120 | 55 | female | 2 | 0 | 0 |
| val75 | 53 | female | 4 | 0 | 0 |
| val106 | 59 | male | 2 | 0 | 0 |
| val109 | 76 | male | 3 | 0 | 0 |
| val64 | 26 | male | 1 | 0 | 0 |
| val65 | 57 | male | 2 | 0 | 0 |
| val97 | 79 | male | 3 | 0 | 0 |
| val124 | 56 | female | 2 | 1 | 0 |
| val132 | 41 | male | 2 | 0 | 0 |
| val121 | 72 | male | 2 | 1 | 0 |
| val49 | 37 | female | 2 | 0 | 0 |
| val88 | 37 | female | 2 | 0 | 0 |
| val67 | 60 | male | 1 | 0 | 0 |
| val52 | 60 | male | 2 | 0 | 0 |
| val118 | 27 | female | 1 | 0 | 0 |
| val99 | 26 | female | 2 | 0 | 0 |
| val70 | 67 | male | 3 | 0 | 0 |
| val117 | 59 | female | 2 | 0 | 0 |
| val127 | 47 | male | 2 | 0 | 0 |
| val95 | 46 | female | 3 | 0 | 0 |
| val133 | 39 | female | 1 | 0 | 0 |
| val46 | 71 | male | 2 | 0 | 0 |
| val44 | 64 | male | 2 | 0 | 0 |
| val84 | 70 | male | 2 | 0 | 0 |
| val81 | 22 | male | 2 | 1 | 0 |
| val111 | 62 | male | 2 | 1 | 0 |
| val115 | 76 | female | 4 | 1 | 0 |
| val122 | 65 | female | 3 | 1 | 0 |
| val30 | 59 | male | 3 | 0 | 0 |
| val94 | 50 | female | 1 | 0 | 0 |
| val69 | 44 | male | 3 | 1 | 0 |
| val18 | 50 | female | 2 | 0 | 0 |
| val63 | 53 | female | 3 | 0 | 0 |
| val119 | 60 | male | 2 | 0 | 0 |
| val126 | 64 | male | 3 | 1 | 0 |
| val93 | 54 | female | 2 | 0 | 0 |
| val103 | 52 | female | 2 | 0 | 0 |
| val76 | 61 | male | 3 | 0 | 0 |
| val116 | 26 | female | 1 | 0 | 0 |
| val91 | 59 | male | 3 | 0 | 0 |
| val112 | 45 | female | 2 | 0 | 0 |
| val21 | 64 | male | 1 | 0 | 0 |
| val128 | 56 | male | 2 | 0 | 0 |
| val50 | 32 | male | 2 | 0 | 0 |
| val107 | 67 | male | 2 | 0 | 0 |
| val27 | 37 | male | 1 | 0 | 0 |
| val48 | 59 | male | 2 | 0 | 0 |
| val57 | 58 | female | 2 | 0 | 0 |
| val59 | 65 | male | 2 | 0 | 0 |
| val87 | 63 | female | 3 | 1 | 0 |
| val96 | 73 | female | 4 | 0 | 0 |
| val53 | 54 | female | 2 | 0 | 0 |
| val45 | 57 | female | 3 | 0 | 0 |
| val71 | 56 | male | 2 | 0 | 0 |
| val19 | 60 | male | 1 | 0 | 0 |
| val20 | 56 | female | 2 | 0 | 0 |
| val31 | 54 | male | 3 | 1 | 0 |
| val35 | 73 | male | 2 | 1 | 0 |
| val78 | 42 | male | 2 | 1 | 0 |
| val125 | 63 | male | 2 | 1 | 0 |
| val92 | 38 | male | 2 | 0 | 0 |
| val28 | 37 | male | 2 | 0 | 0 |
| val82 | 72 | male | 2 | 0 | 0 |
| val108 | 77 | male | 3 | 0 | 0 |
| val90 | 63 | female | 4 | 1 | 0 |
| val47 | 51 | female | 2 | 0 | 0 |
| val113 | 45 | female | 1 | 0 | 0 |
| val98 | 51 | male | 4 | 0 | 0 |
| val79 | 26 | female | 1 | 1 | 0 |
| val56 | 78 | female | 4 | 0 | 0 |
| val104 | 69 | male | 3 | 1 | 0 |
| val105 | 60 | male | 2 | 0 | 0 |
| val89 | 72 | female | 4 | 1 | 0 |
| val62 | 61 | male | 4 | 0 | 0 |
| val36 | 74 | male | 2 | 1 | 0 |
| val74 | 49 | female | 2 | 0 | 0 |
| val22 | 36 | female | 2 | 0 | 0 |
| val129 | 71 | male | 1 | 0 | 0 |
| val60 | 46 | female | 3 | 0 | 0 |
| val114 | 74 | female | 3 | 0 | 0 |

## Model Summaries

### Summary of the model assessing the influence of pathology results on IG clonality in the pre-ICI era cohort

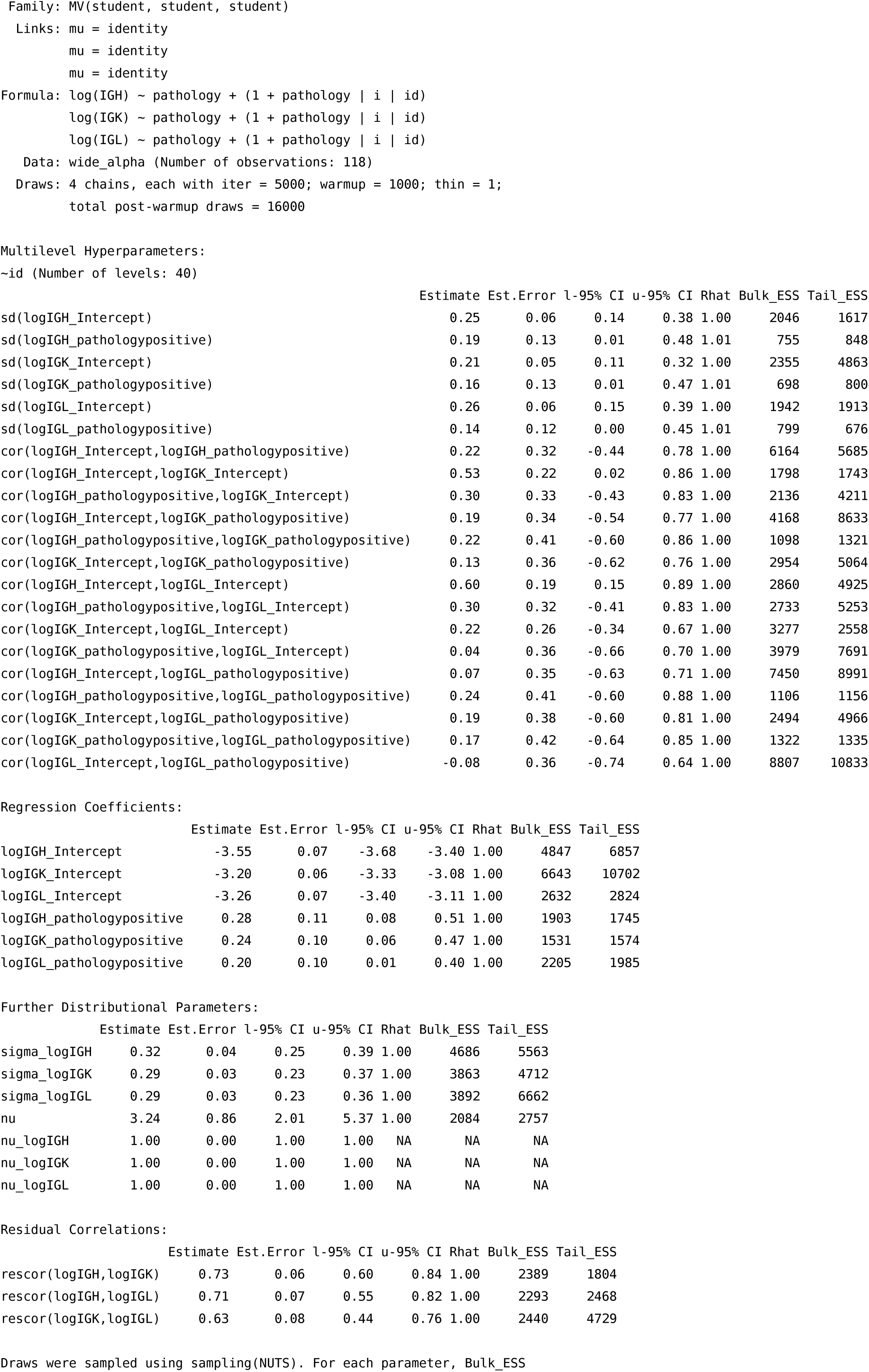
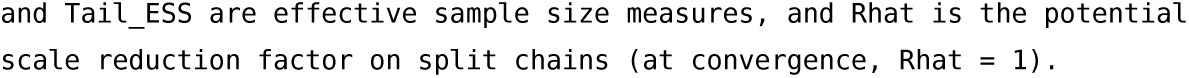

### Summary of the model assessing the influence of pathology results on TCR clonality in the pre-ICI era cohort

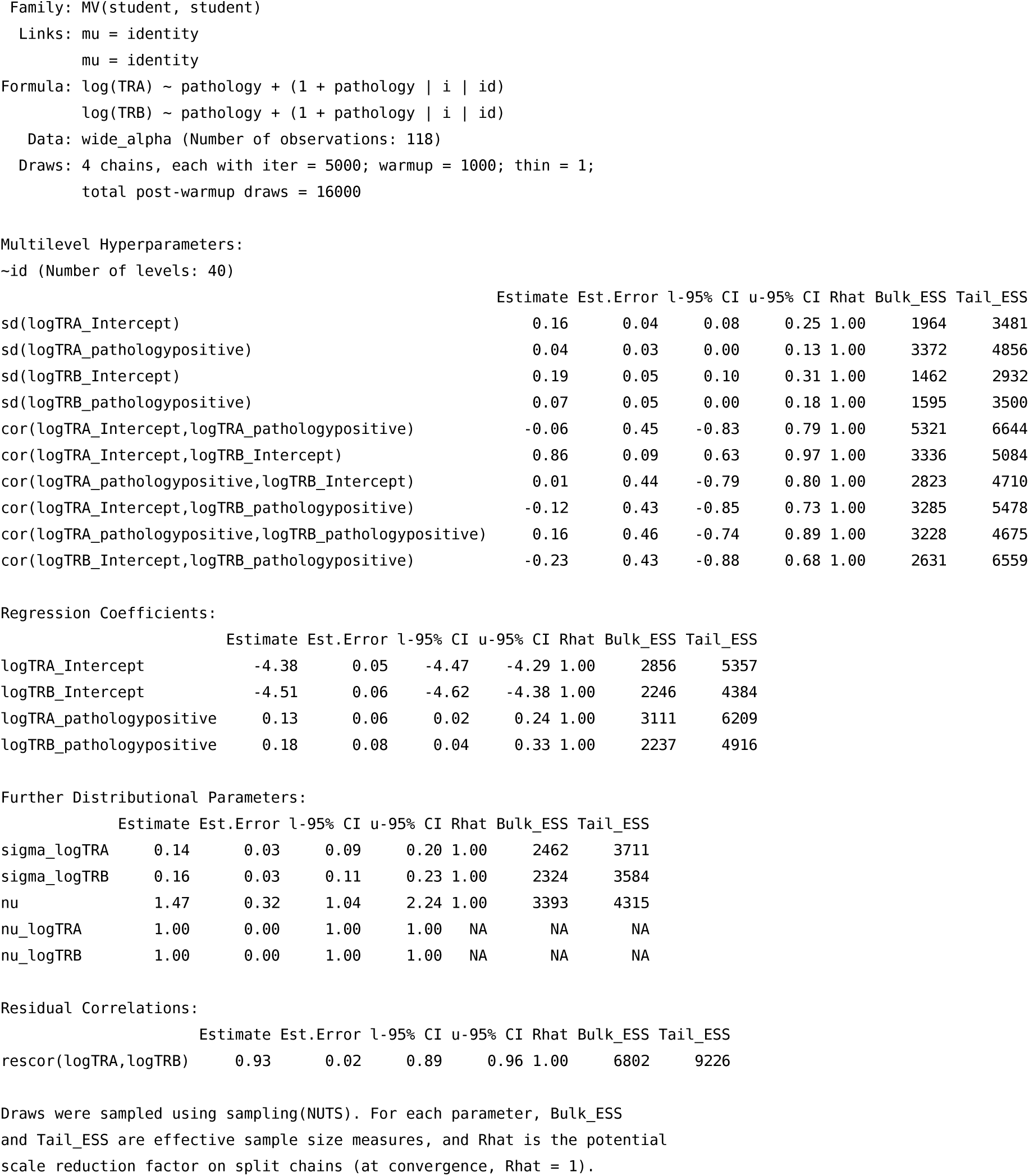

### Summary of the model assessing the influence of the T stage on IG clonality in the pre-ICI era cohort

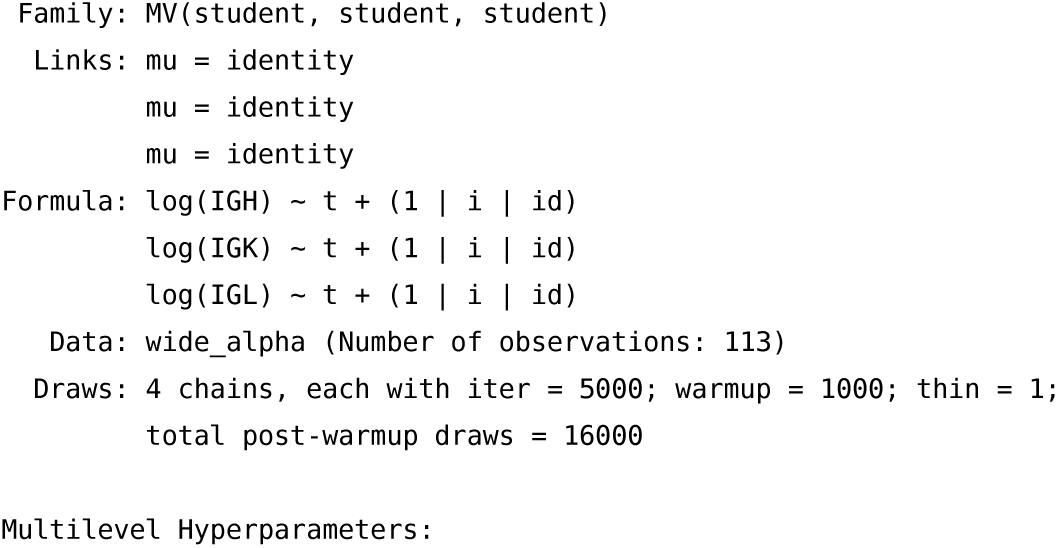
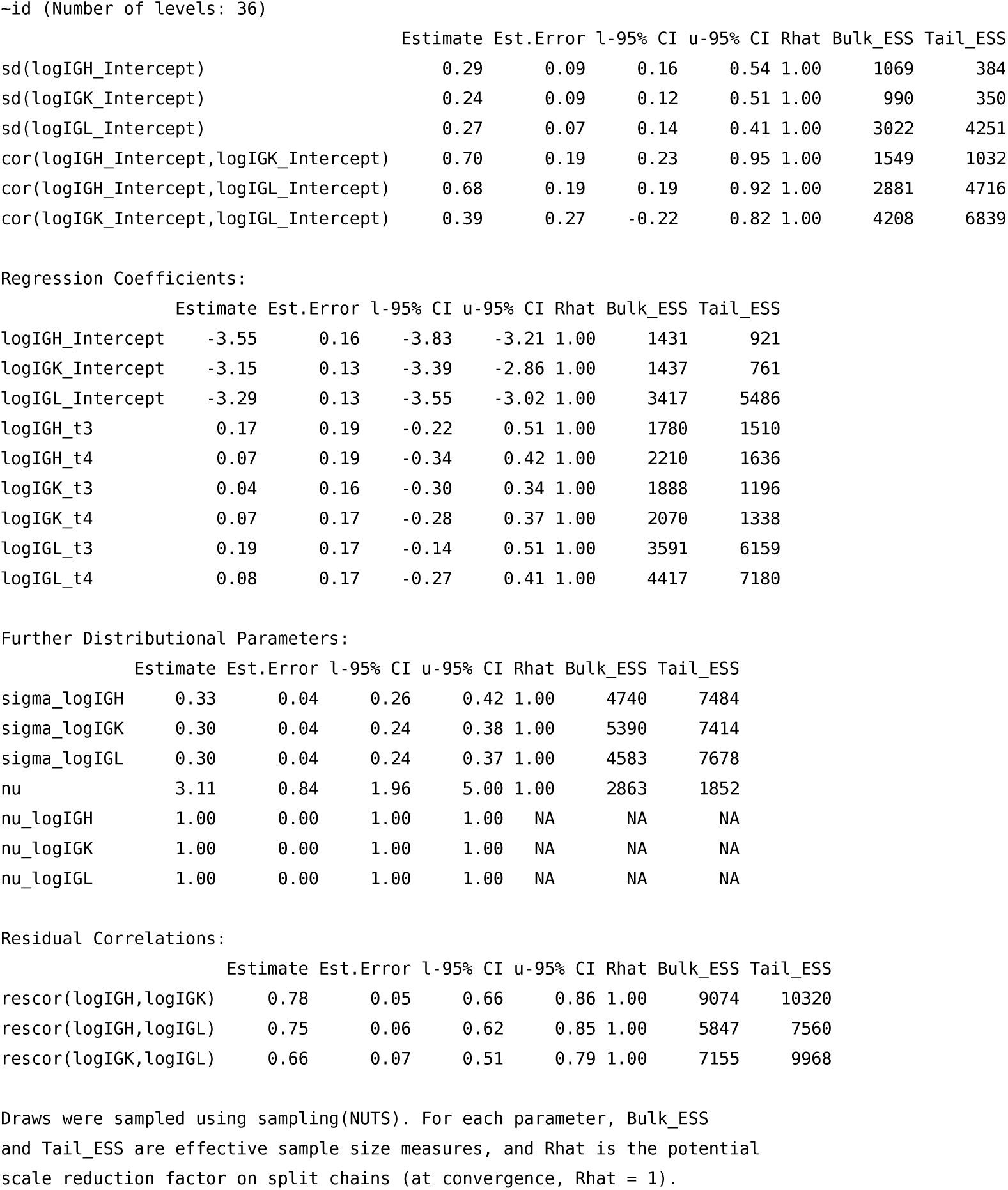

### Summary of the model assessing the influence of the T stage results on TCR clonality in the pre-ICI era cohort

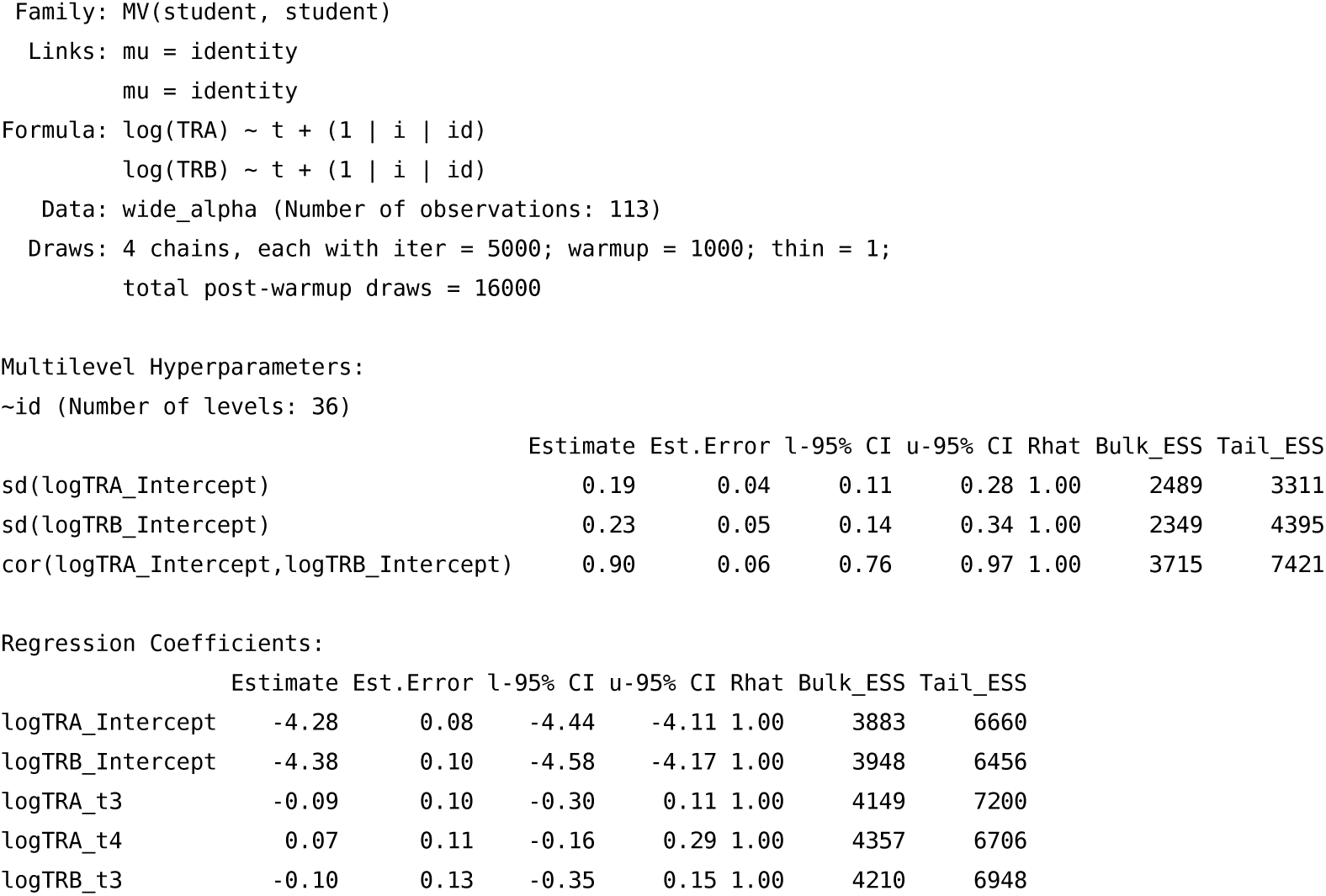
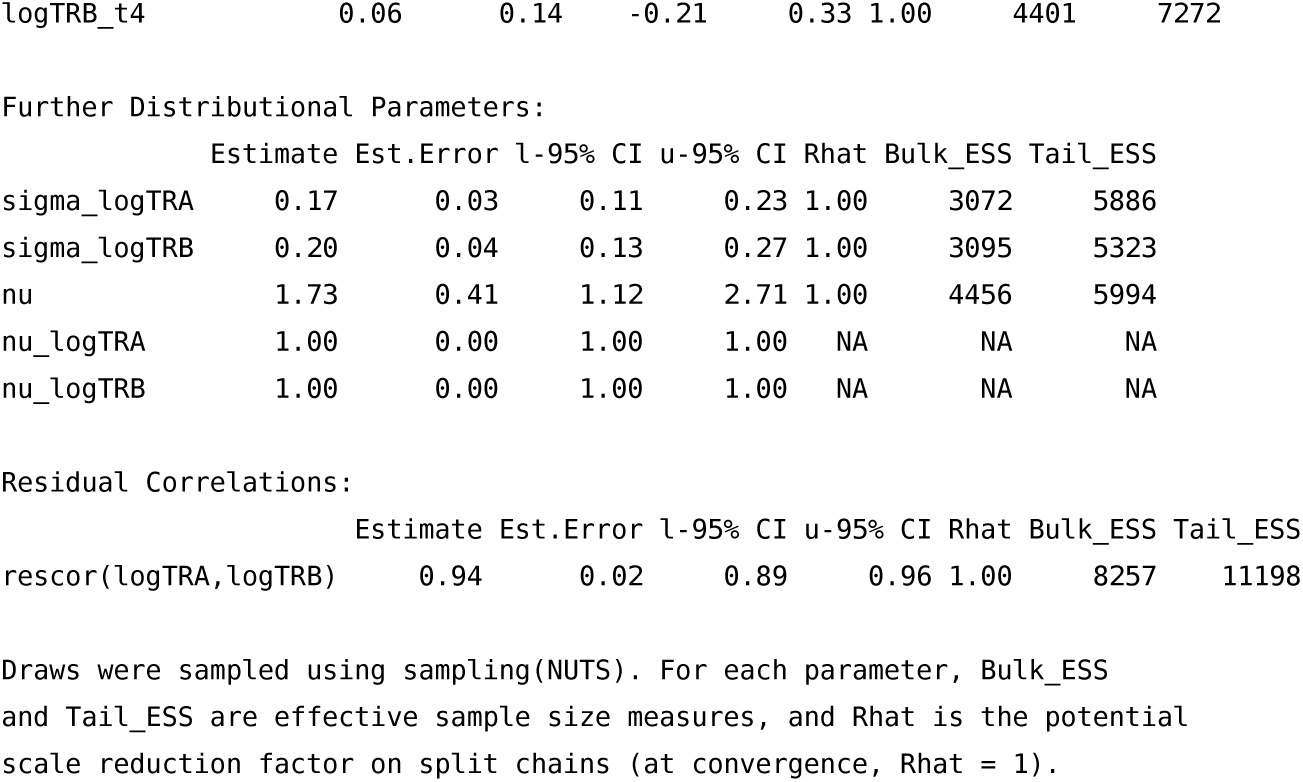

### Summary of the model assessing the influence of the N stage on IG clonality in the pre-ICI era cohort

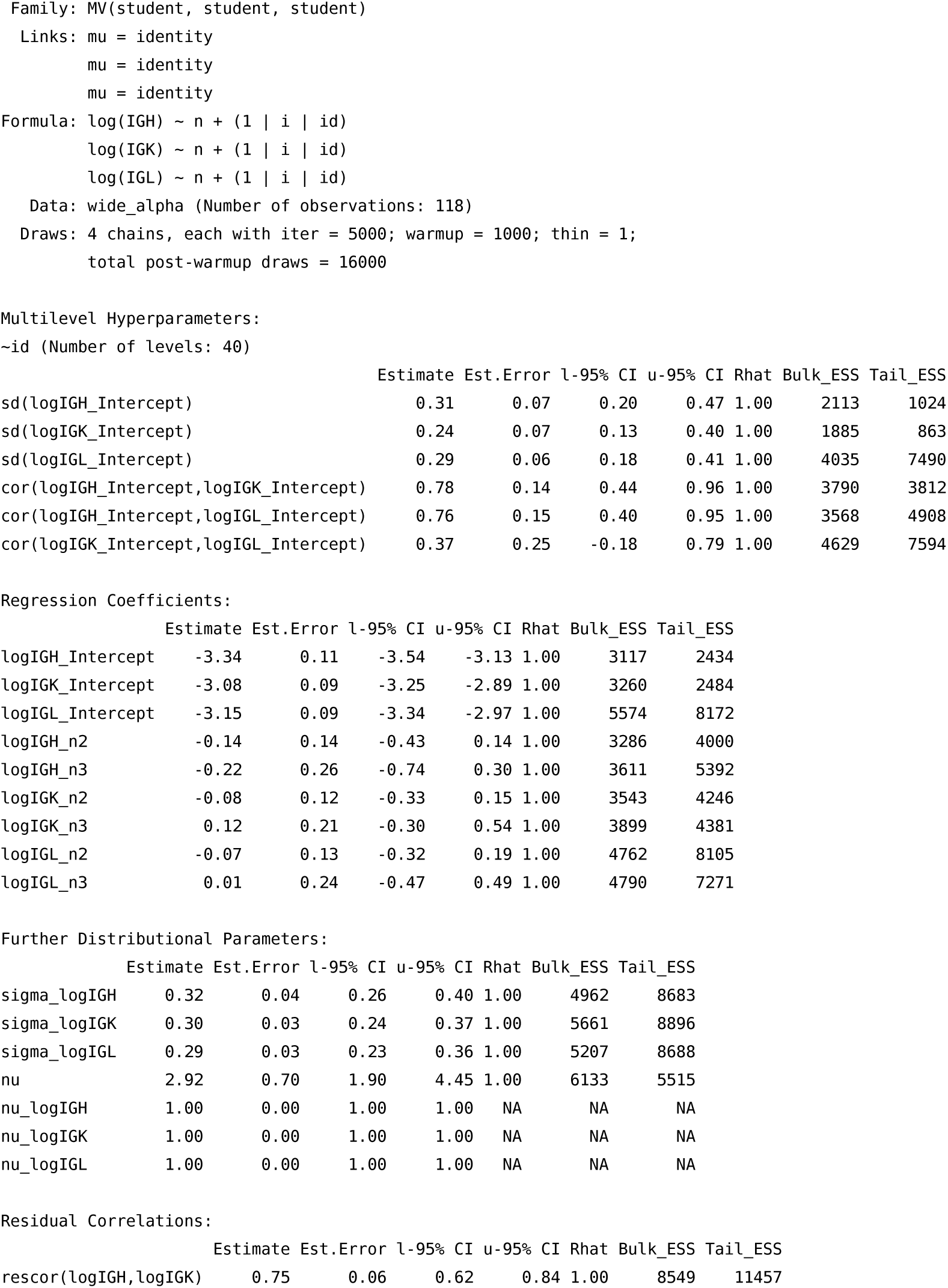
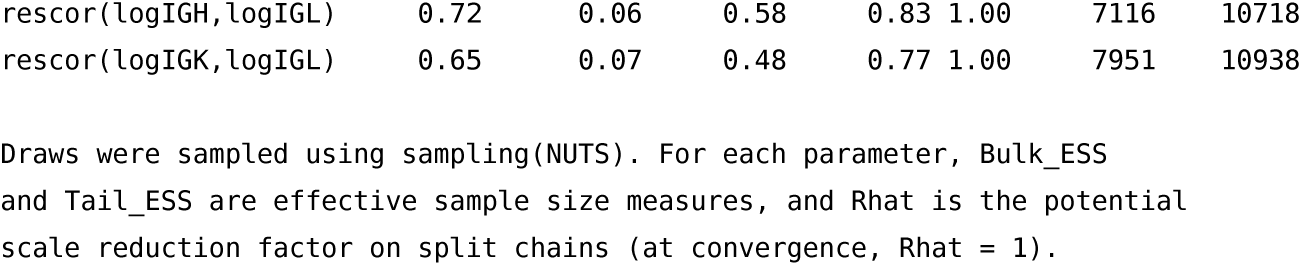

### Summary of the model assessing the influence of the N stage results on TCR clonality in the pre-ICI era cohort

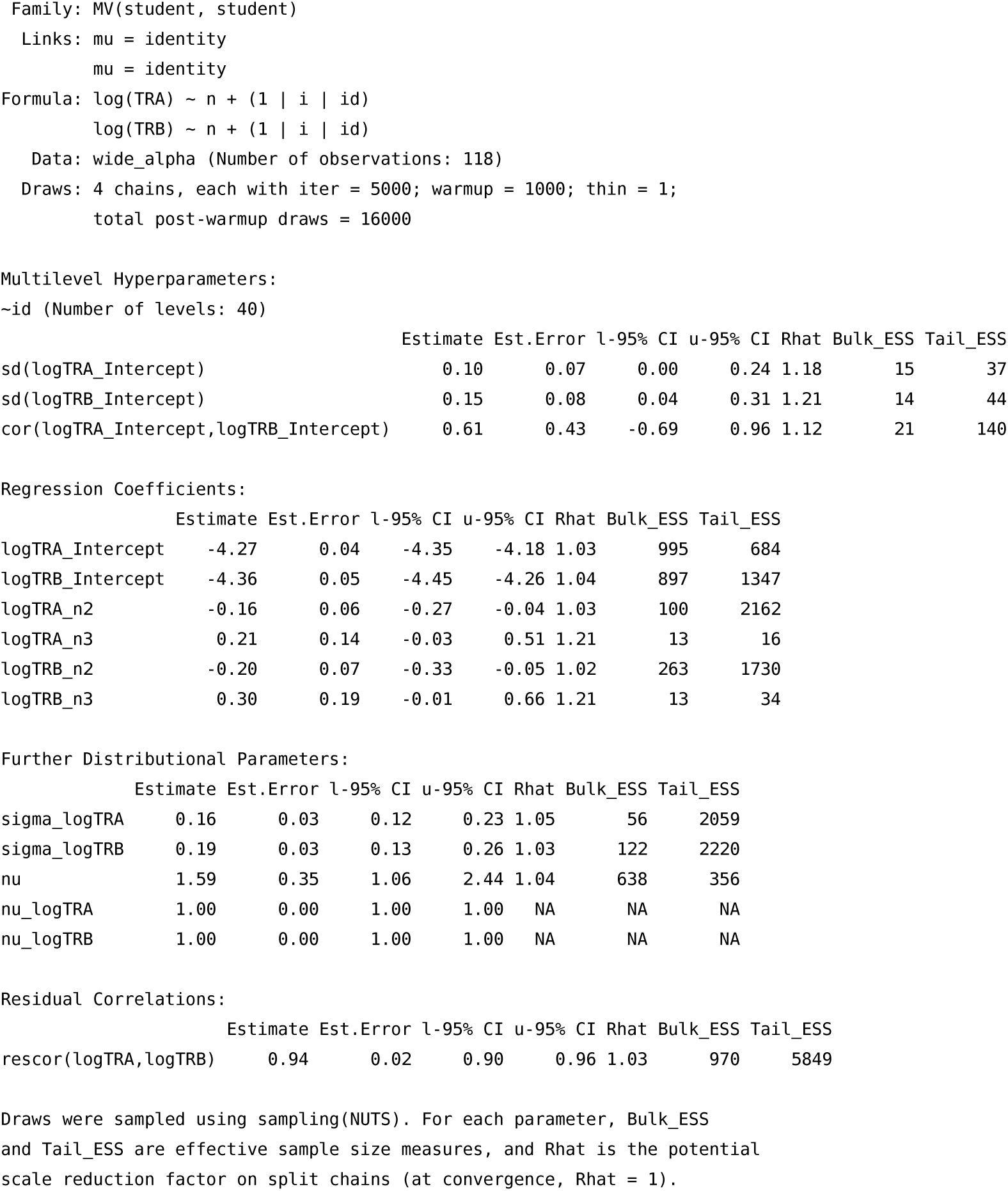

### Summary of the model assessing the influence of the UICC stage on IG clonality in the pre-ICI era cohort

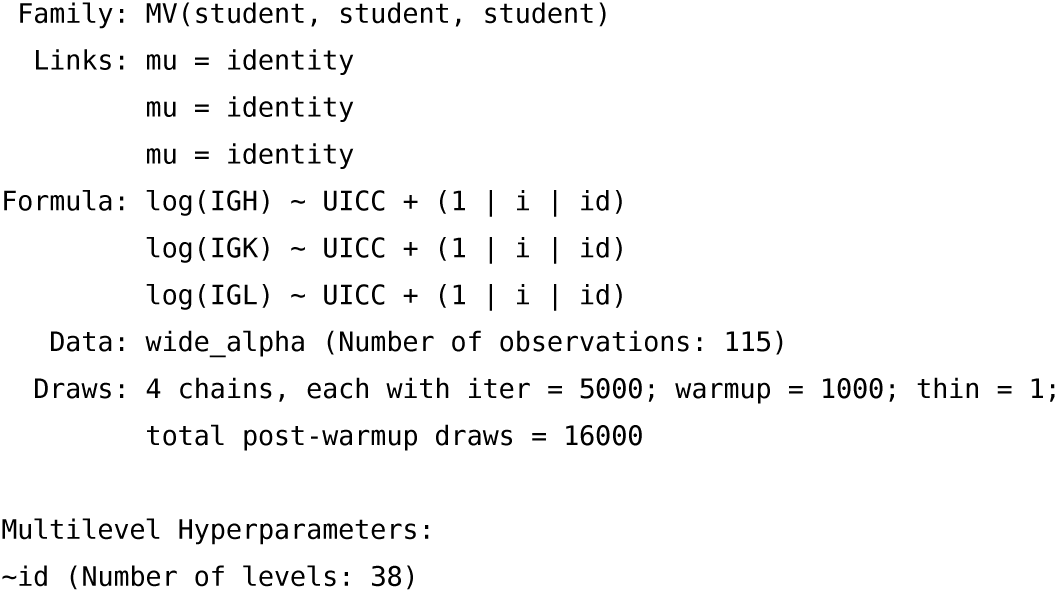
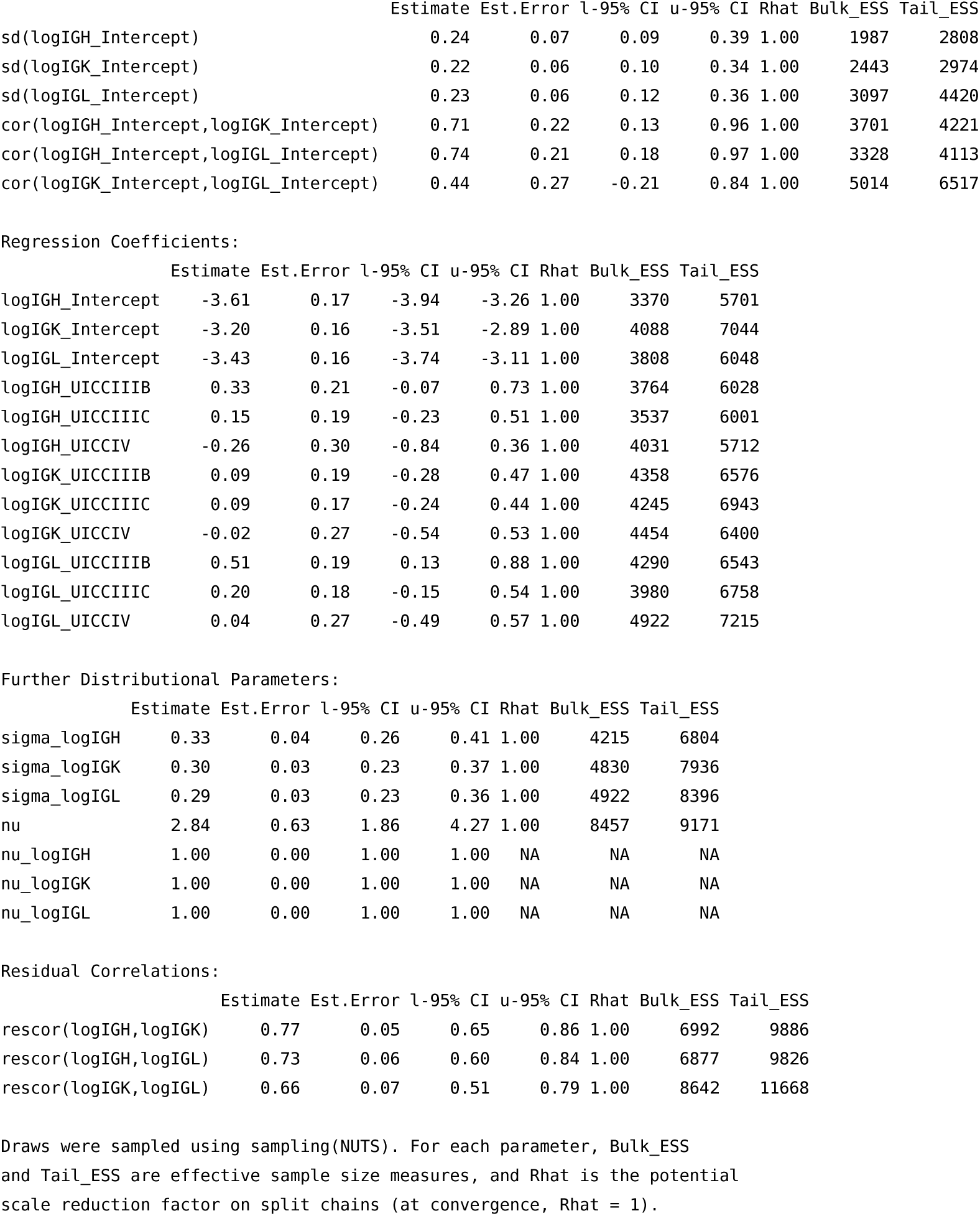

### Summary of the model assessing the influence of the UICC stage results on TCR clonality in the pre-ICI era cohort

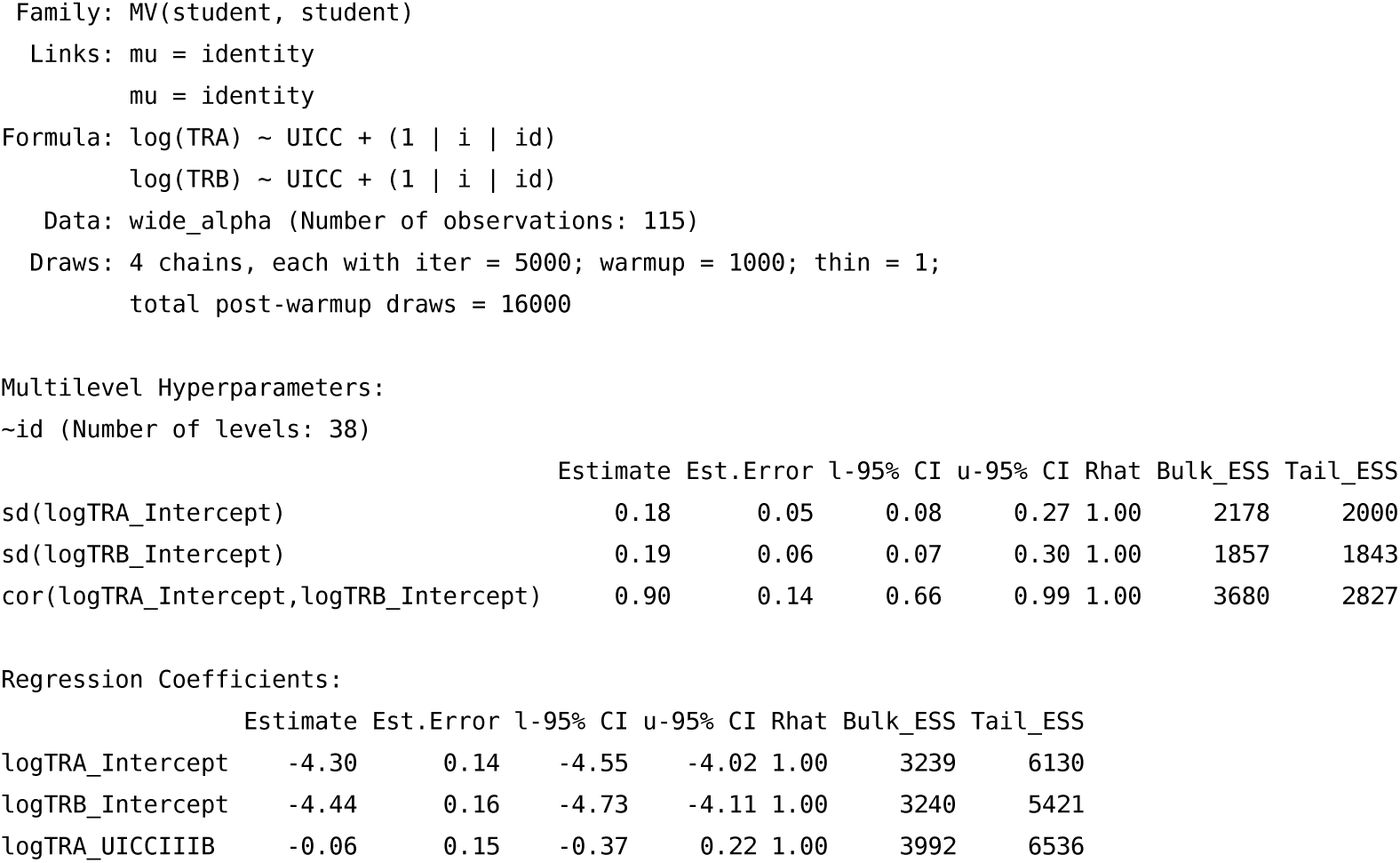
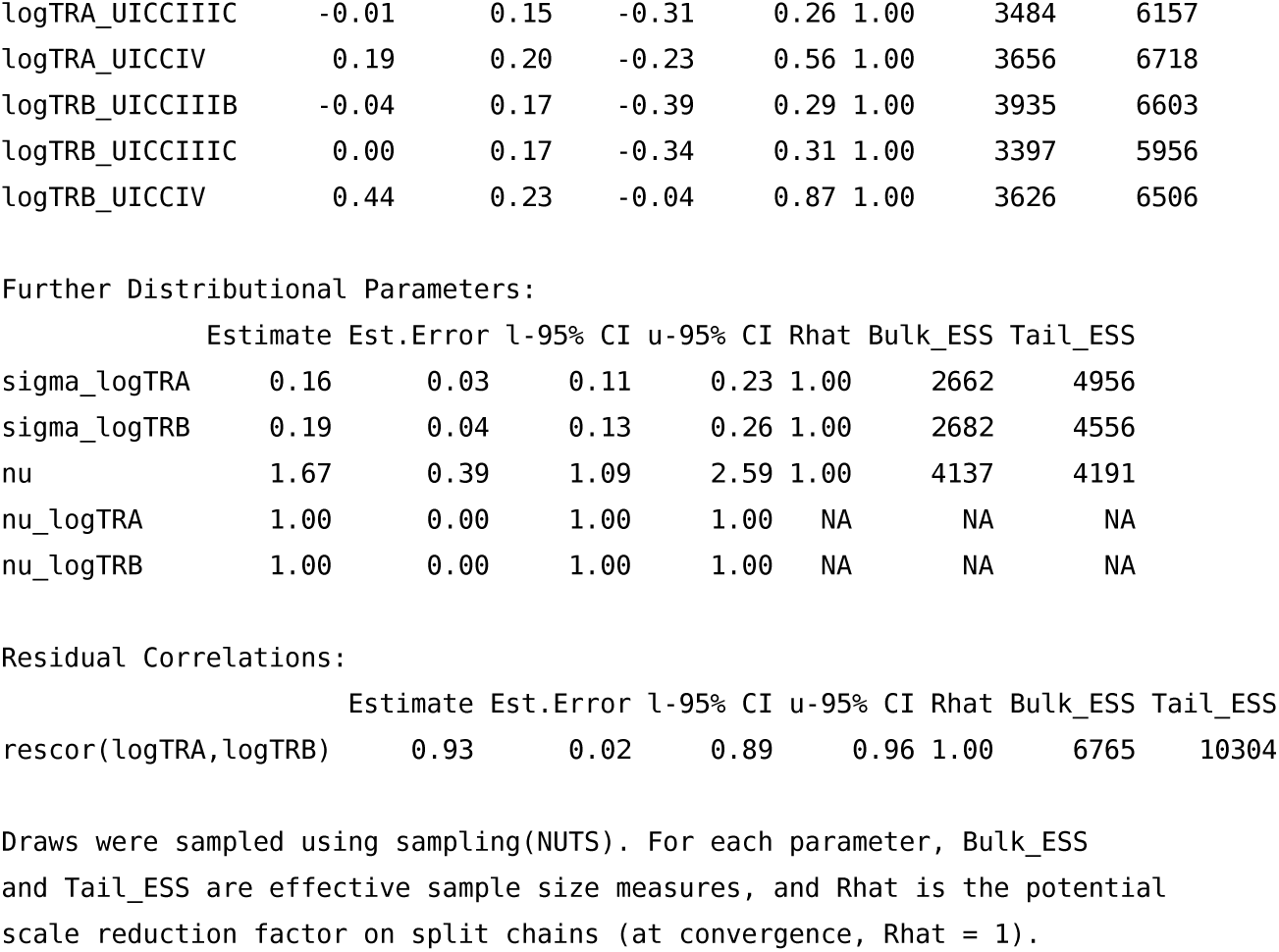

### Summary of the model assessing the influence of IG clonality on survivorship in the pre-ICI era cohort

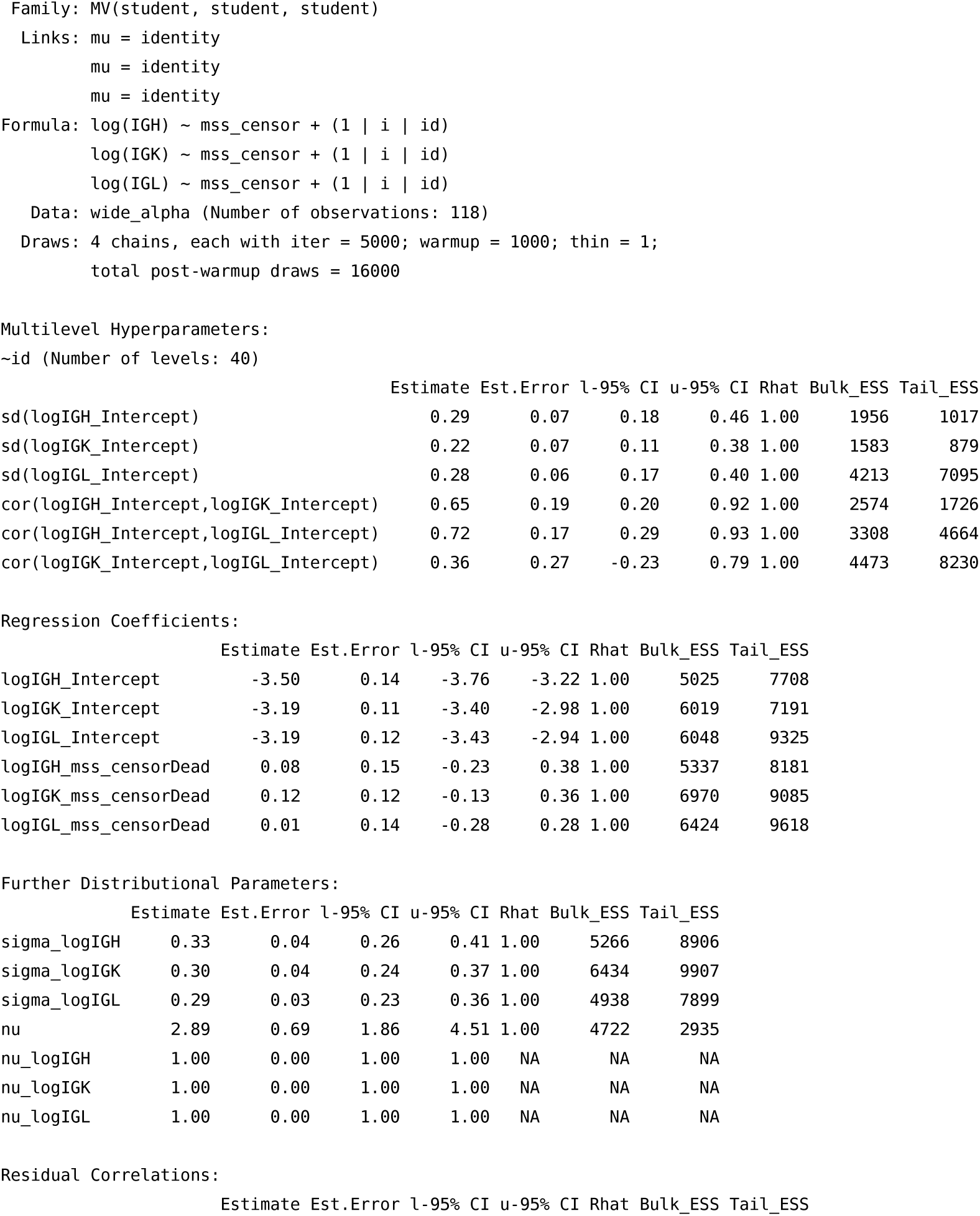
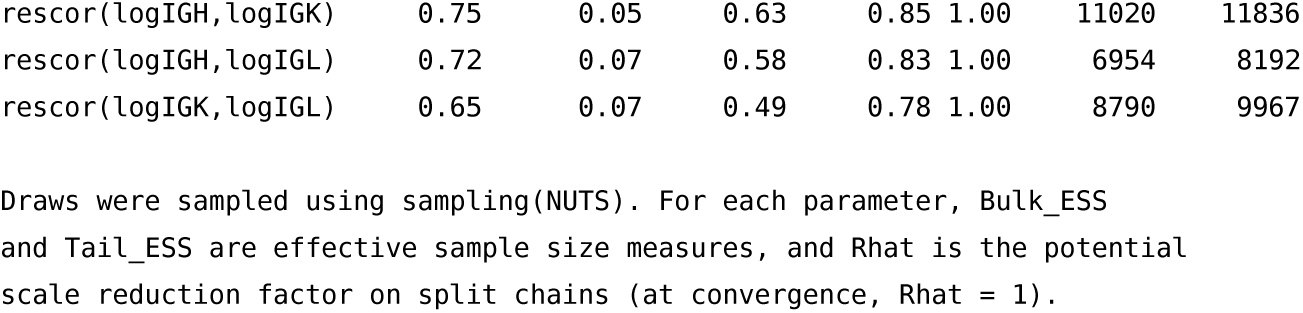

### Summary of the model assessing the influence of TCR clonality on survivorship in the pre-ICI era cohort

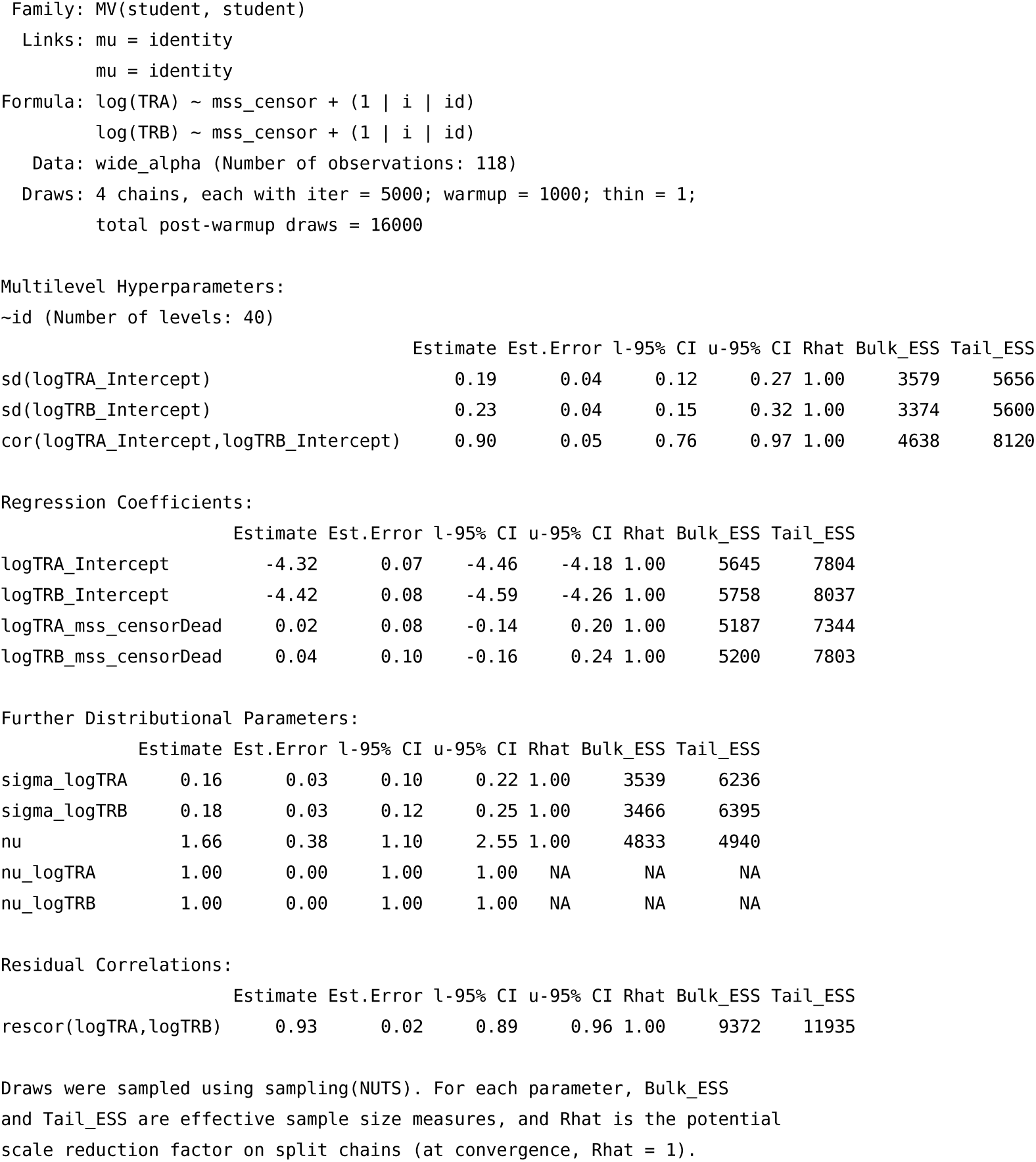

### Summary of the model assessing the influence of pathology results on LN IG clonality in the contemporary validation LN cohort

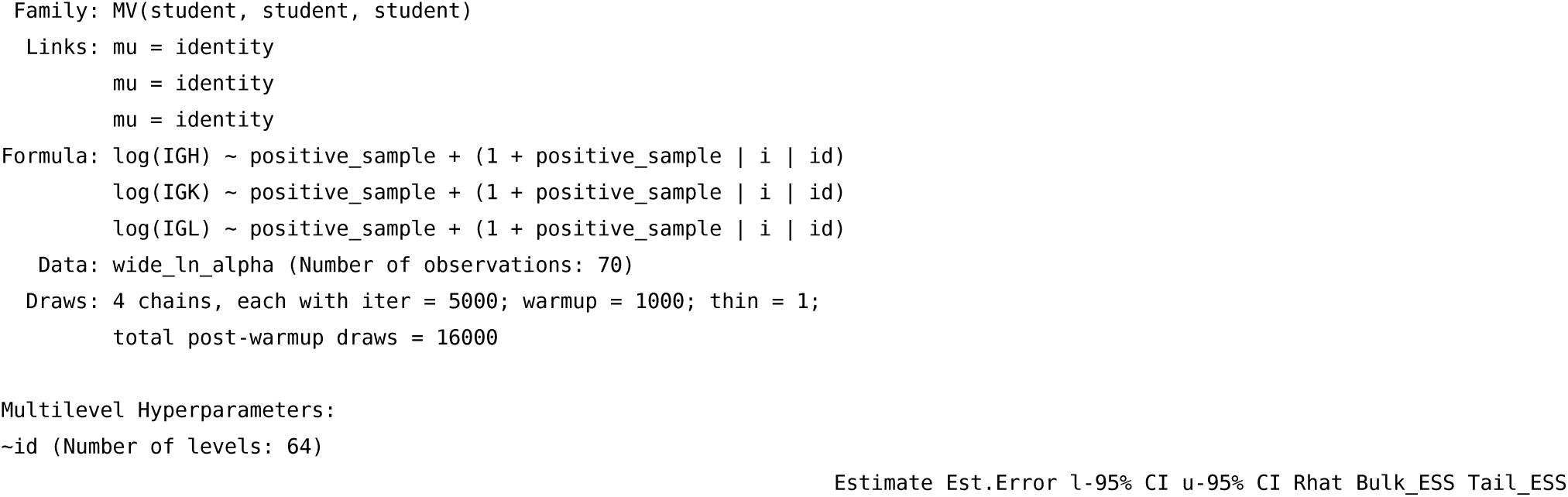
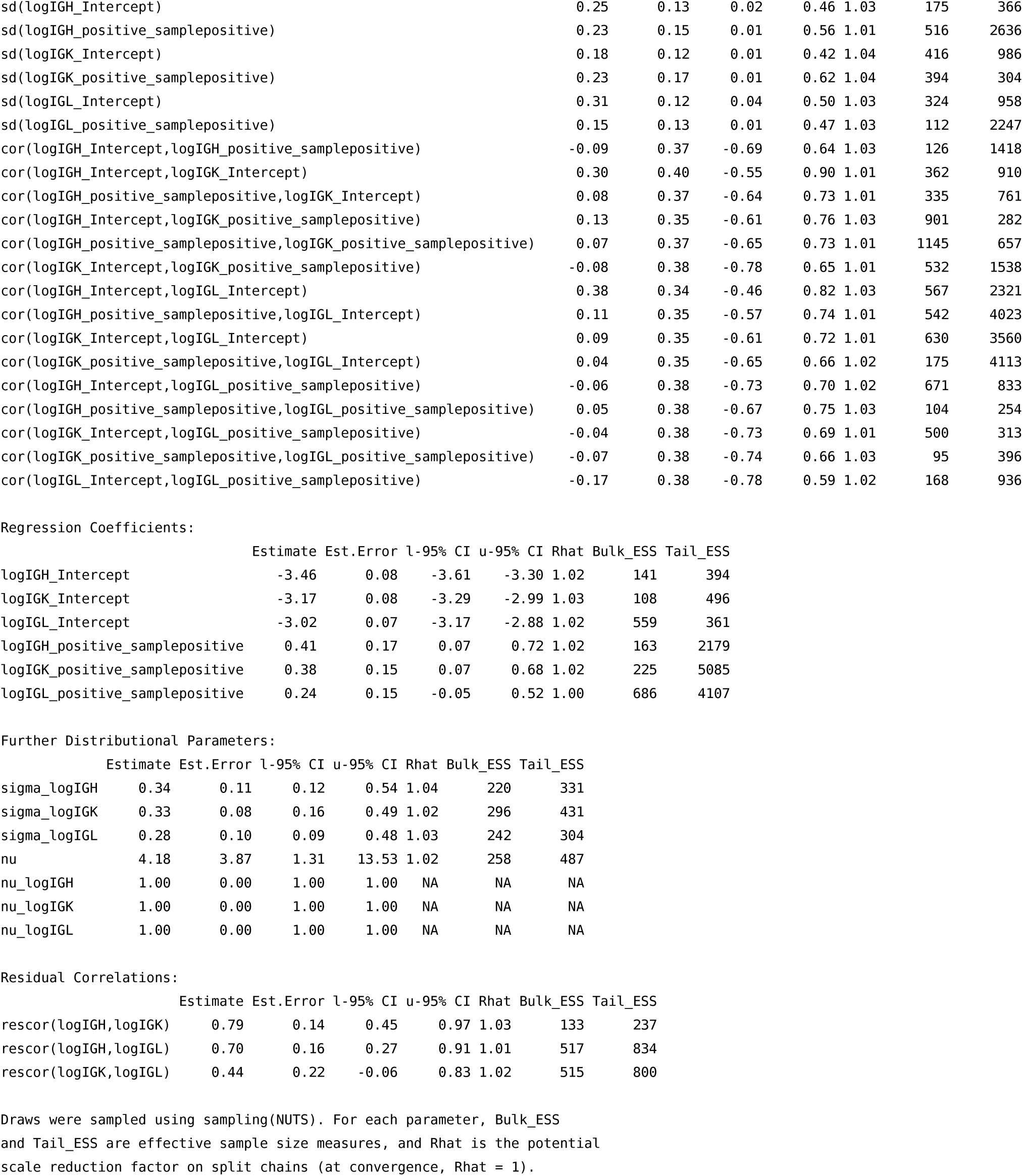

### Summary of the model assessing the influence of pathology results on LN TCR clonality in the contemporary validation LN cohort

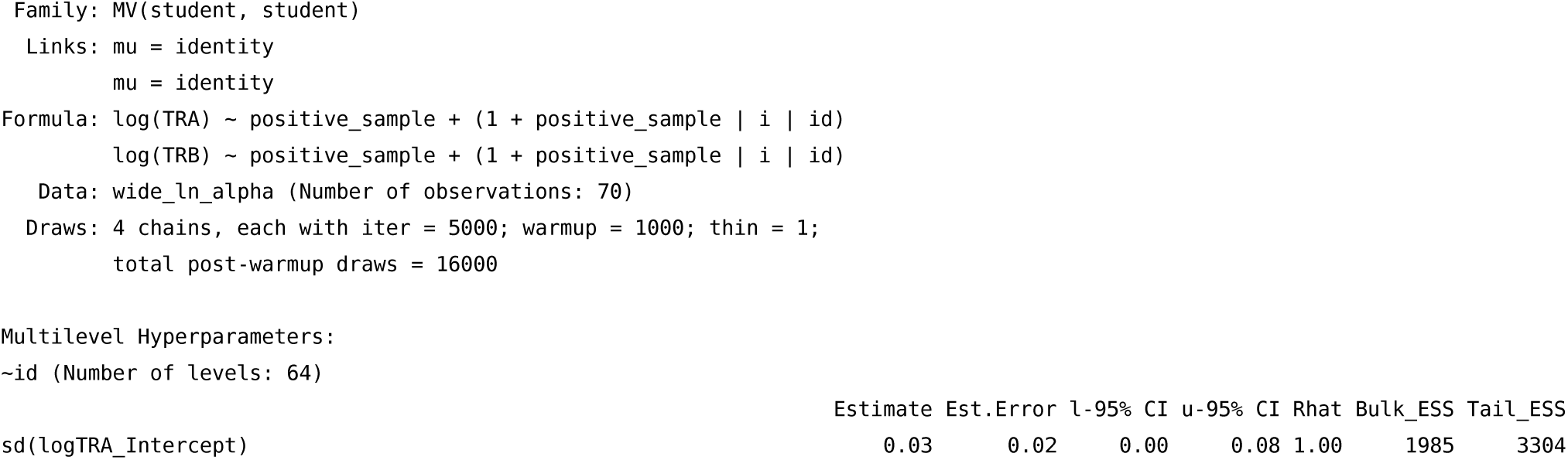
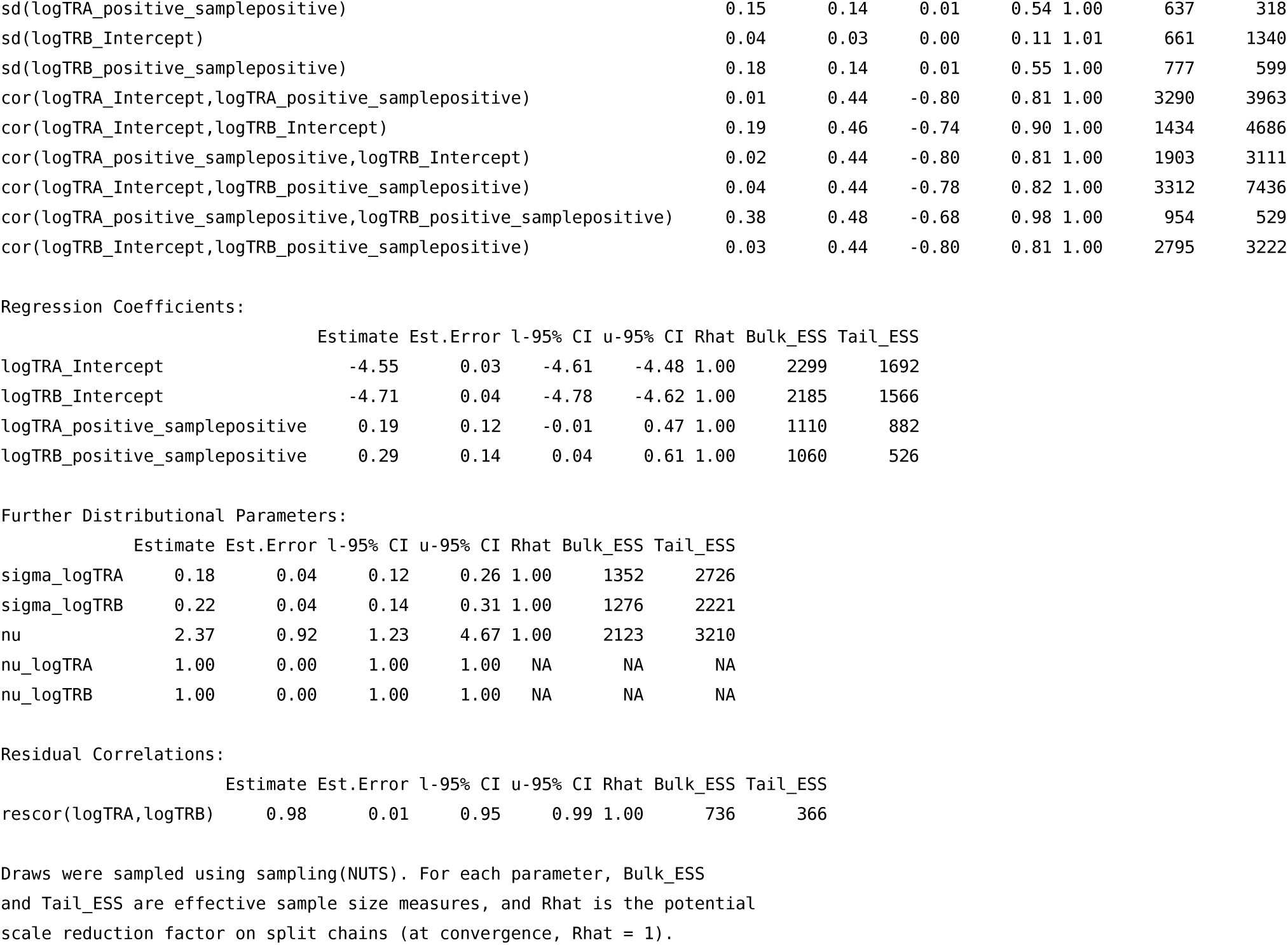

### Summary of the model assessing the influence of pathology results on PBMC IG clonality in the contemporary validation LN cohort

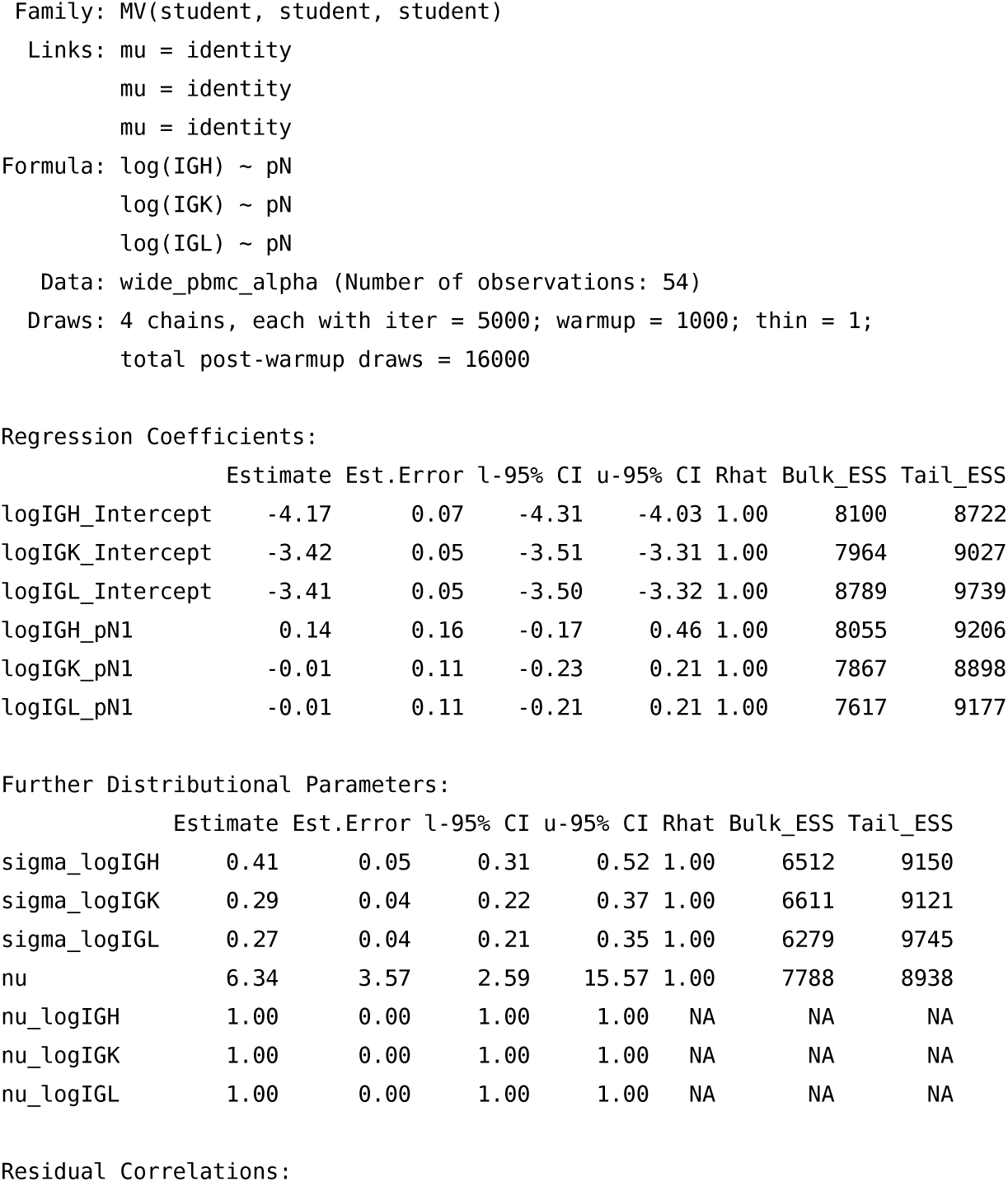
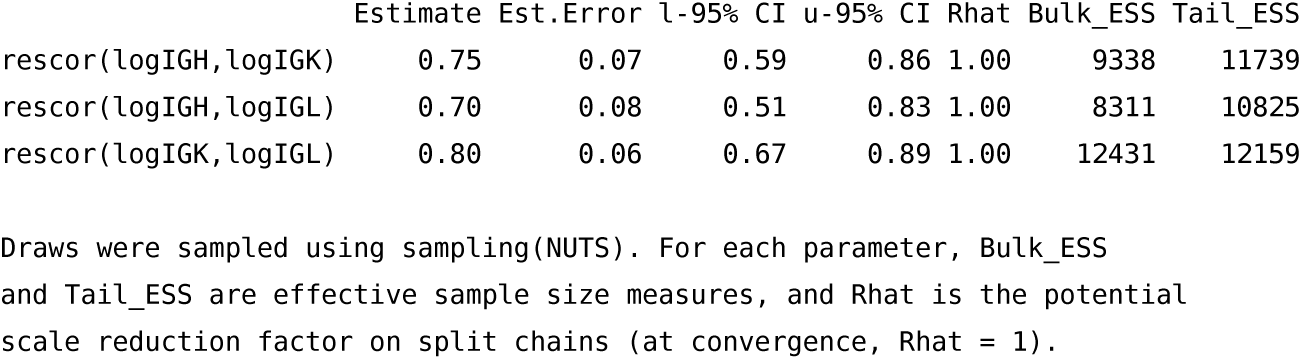

### Summary of the model assessing the influence of pathology results on PBMC TCR clonality in the contemporary validation LN cohort

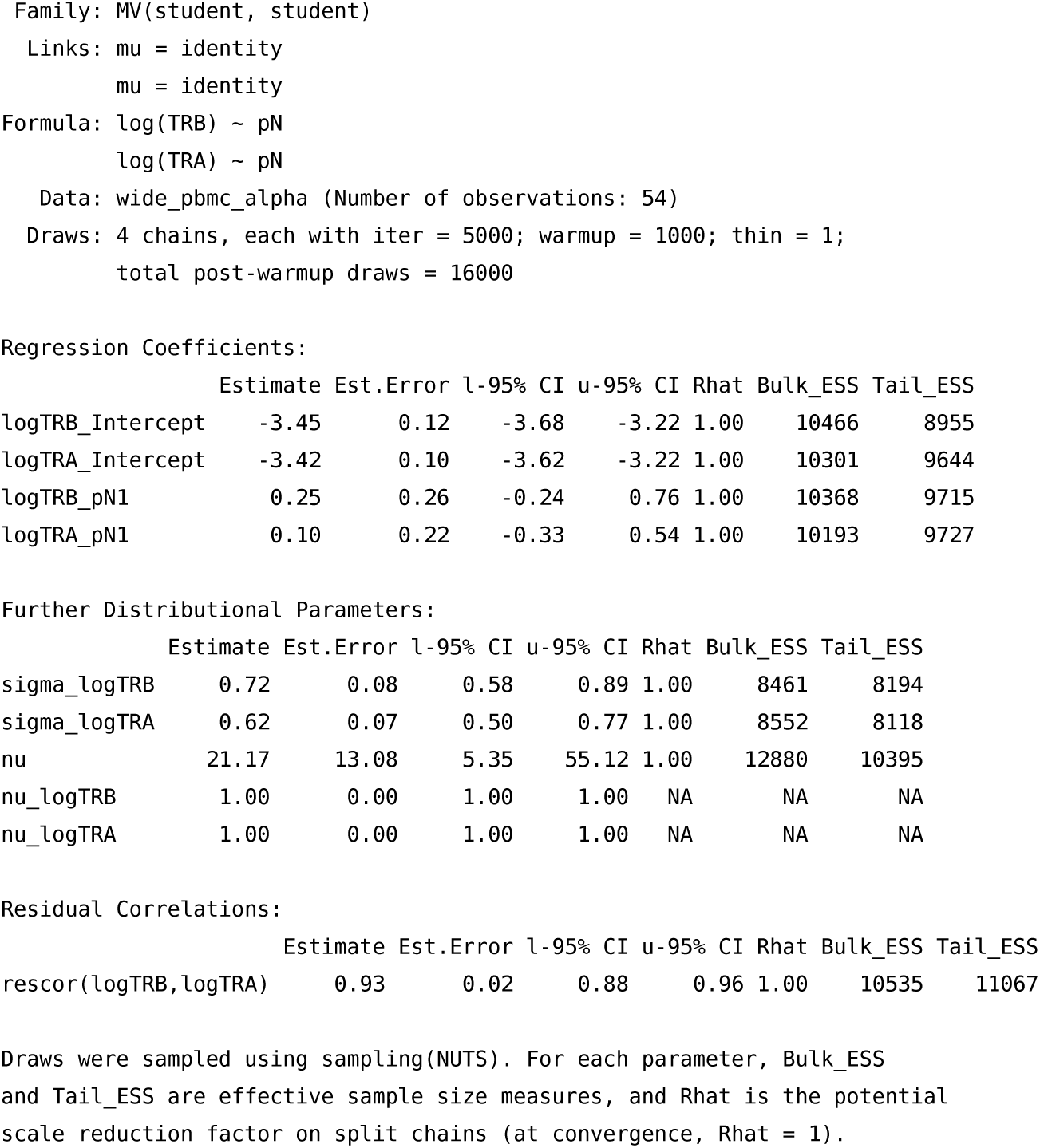

